# A metabolic perspective on polyploid invasion and the emergence of life histories: insights from a mechanistic model

**DOI:** 10.1101/2023.09.11.557145

**Authors:** Silvija Milosavljevic, Felipe Kauai, Frederik Mortier, Yves Van de Peer, Dries Bonte

## Abstract

Whole genome duplication (WGD, polyploidization), the fusion of unreduced gametes, has been identified as a driver of genetic and phenotypic novelty. Unreduced gamete formation is common in a wide range of species, but surprisingly, few polyploidization events have shown to be ecologically successful. Positive density dependence, by minority cytotype exclusion, and niche shifts are currently considered the most important drivers behind ecological failure or success. Genome doubling also results in increased cell sizes and metabolic expenses which, on their own may be sufficient to drive polyploid establishment in stable environments where their simple ancestors thrive.

We developed a mechanistic model, motivated by data from natural plant polyploid species, to test whether realistic changes in size and metabolic efficiency allow polyploids to coexist with, or even invade, their original diploid population. Central to the model is metabolic efficiency, a functional trait that determines how energy gained from size-dependent photosynthetic metabolism is allocated to basal metabolism, somatic growth and reproductive growth.

Polyploid invasion was observed across a wide range of metabolic efficiency differences between polyploids and their ancestors. Higher metabolic efficiency facilitates polyploid invasion, but even with minor deficits, establishment was facilitated by recurrent formation in these settings of high competition for nutrients. Interestingly, a long-term coexistence with the diploid ancestor was found to be possible only within a narrow range of this parameter space. Perenniality of the plants did not qualitatively affect these insights. Feedbacks between size-dependent metabolism and allocation of gained energy generated eventually size and age differences, resulting in intra- and intercytotype competition for nutrients as the major force for population dynamics. We thus demonstrate that changes in metabolic efficiency on their own are sufficient to impose establishment, but these advantages do not need to be substantial.

## Introduction

Polyploids are organisms that gained a whole set of chromosomes through whole genome duplication (WGD), usually due to unreduced gametes fusion (Van de Peer et al. 2017). Polyploidization is considered important for genome evolution and is a common mode of speciation. Polyploids can be found in all domains of life, including animals (Mable et al. 2011, Otto & Whitton 2000), fungi (Albertin and Marullo 2012, Campbell et al. 2016, Todd et al. 2017), bacteria and archaea (Breuert et al. 2006, Pecoraro et al. 2011), but most commonly found and studied in plants.

Ancient genome duplications are rare and appear to have established during specific periods in evolution, suggesting that stressful environmental conditions facilitate polyploid success (Fawcett et al. 2009; Vanneste et al. 2014a; Vanneste et al. 2014b, Lohaus & Van de Peer, 2016; Cannon et al. 2015; Yu et al. 2017; Huang et al. 2016). At contemporary time scales, the widespread presence of polyploids has equally been linked to their increased stress tolerance (Chao et al. 2013, Yang et al. 2014, Marks et al. 2023), but also to acquired niche shifts from changes in size, floral traits and life history (Ramsey & Ramsey 2014, Kiedrzyński et al 2021). Adaptations leading to niche shifts, selfing and asexual reproduction (Novikova et al. 2022) mitigate the effect of minority cytotype exclusion (MCE), a process of positive density dependence hindering successful mating and reproduction at low density due to incompatibilities between the simple and doubled genotypes (Levin 1975). Also perenniality is considered to be advantageous as it provides a longer time-window for reproduction of the newly formed polyploid thus reducing MCE (Gustafsson 1948, Van Drunen & Friedman 2022).

Genotypic and phenotypic changes associated with an increase in cell, organ and body size, impose serious costs by modifications of the cellular management, energy use, cell division abnormalities, and genomic instability (Ramsey & Schemske 2002, Van Drunen & Husband 2019). Increases in size, and associated changes in energy use and metabolism, as expressed in photosynthesis and gas exchange are among most commonly reported changes after WGD (Bomblies 2020). Typically, metabolic rate increases with body size, but this increase lowers with increased body size. This allometric scaling relationship between metabolic rate and body mass is fundamental to biology, but also one of the most debated concepts in ecology and evolution (White et al. 2007, Isaac & Carbone 2010). It makes size of an individual key to its behavior and physiology, as it sets the ecophysiological boundaries by mediating energy intake, use and thus efficiency (Peters 1983). Body size thus not only affects individual-level metabolic processes but also higher ecosystem functioning (Yvon-Durocher & Allen 2012) through its impact on life histories and the resulting population dynamics (Price et al. 2010). The exact impact of metabolic processes on life history will depend on how energy is used and allocated among vital functions. This allocation is central to the Dynamic Energy Budget (DEB) theory (Kooijman 2010).

To understand how polyploidy induced changes in metabolism affect polyploid establishment through changes in size and life history, we present a new mechanistic model in which metabolic and reproductive activity – conditional to body size and resource availability – are mediated by the kinetics of photosynthetic rate (Hu et al 2021). We examine how changing metabolic efficiency in emerging polyploids affects their chances of establishment while competing with diploids. To establish, a neopolyploid must demonstrate positive (sub)population growth while interacting with its environment, typically where the original cytotype is already established. One crucial requirement for the neopolyploid’s successful invasion is to possess a higher or at least comparable average fitness level. However, since polyploidization increases cell size, leading to longer cell cycles, reduced metabolic efficiency and a slower growth (Bennett 1972), it is assumed that adaptations to offset that cost are necessary. Our hypothesis is that neopolyploids must possess at least the same metabolic efficiency as diploids to establish and invade successfully, or adapt their life history to offset metabolic inefficiency. Demographic effects of different life history strategies may evolve to counterbalance these costs, such as the emergence of perenniality and a later maturation age, which could aid in reproductive assurance (Van Drunen & Husband 2019). Therefore, we also model transitions from annuals to perennials.

This study represents the first investigation into eco-evolutionary dynamics in polyploid establishment resulting from basic ecological mechanisms driven by documented changes in metabolism. We believe that taking this mechanistic modeling approach with ecology and physiology in mind is a necessary interdisciplinary way of addressing longstanding questions about polyploid establishment (Soltis et al. 2016). While ecophysiological insights have so far not been applied in polyploidy research, these approaches are opening new avenues and changing our views in other ecological fields. For example, using biophysical mechanistic modelling, Strubbe et al. (2023) showed that potential ranges of invasive bird species are constrained by body size and metabolic rates, forecasting a highly likely further spread of these non-native species. Using DEB models has also been proven useful for improving ecological predictions in variable environments (Monaco & McQuaid 2018), especially in ecotoxicology, where this type of models aid in transforming individual level responses and traits into population and higher-level dynamics (Jager et al. 2014). This research approach shifts the focus from population genetics models to more mechanistic ecological models, shedding light on the fundamental factors influencing polyploid success in ecological contexts.

## Materials and methods

We developed a consumers-nutrients model to investigate the conditions under which polyploids can coexist or invade an initially diploid population when metabolism and reproduction depend on body size. The model is a spatially explicit, discrete-time model where one time step corresponds to one day in the lifetime of consumer individuals and 100 days correspond to one year or growing season. This mechanistic model is based on theory of polyploid formation and phenotypic changes. We base the model on diploid plant population with emerging tetraploids and parametrize the photosynthetic rate with non-woody plants model fits from Hu et al. (2021). By applying an individual-based approach, we can incorporate intraspecific size variation and stochasticity during the initialization of individuals within the model. No explicit sexual reproduction is modelled here, so it can be interpreted as a model of asexually reproducing and clonal organisms, but also as female-only model of sexually reproducing organisms or unicellulars model. Due to this, the model does not include minority cytotype exclusion as such.

We provide detailed description of the model in the Supplementary Material 1, according to the ODD protocol (Overview, Design concepts, Details) for describing individual-based models (Grimm et al. 2006, 2010, 2020). An overview of all parameters is available in the Supplementary Material 1 as well.

### The landscape

The landscape is a lattice of 40x40 cells, where each cell has certain amount of resources (nutrients), represented as mass in grams. The edges of the landscape are not wrapped so there can be effect on the seed dispersal. So model is spatially explicit and discrete.

### The resource

Resources are modelled as each landscape cell’s property and each cell is initialized with 1 g of resources, as an abstraction of main nutrients needed for plant growth like nitrogen and phosphorus. They are supplied with rate r (g/day) from undefined external sources up to the maximum amount of resources Rmax (g).

### The consumers

Within the landscape, each consumer is individually modeled as either diploid or tetraploid. Each day, individuals have the opportunity to uptake nutrients and grow, followed by a chance to reproduce after a season of 100 days (Figure 1).

**Figure 1.**
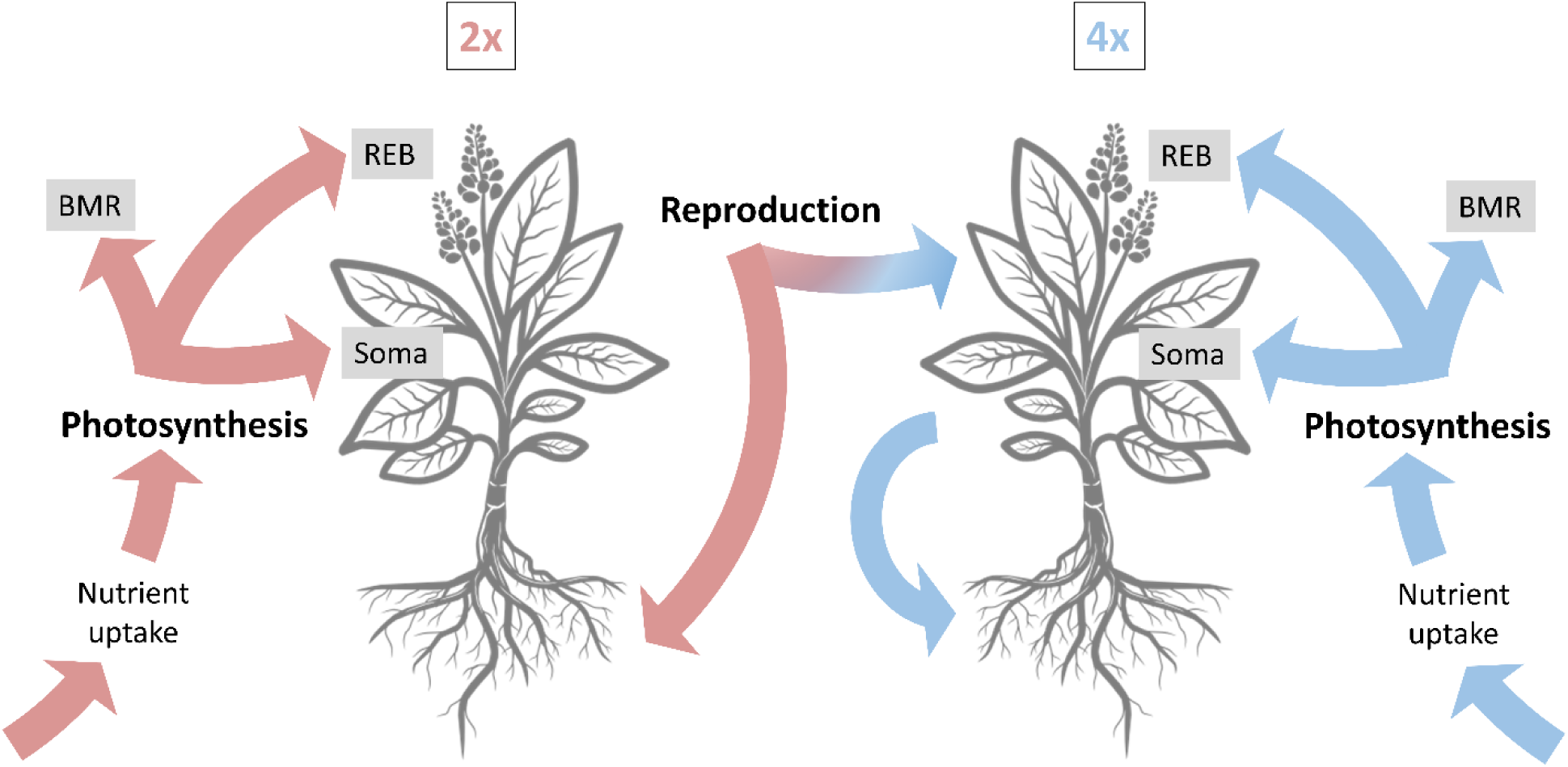
Graphic representation of the processes implemented in the model. All processes happen on daily basis, except reproduction which happens after 100 days (1 season). Red colors are for diploids, and blue for tetraploids throughout the manuscript. Notice that tetraploid individuals only produce tetraploid offspring. BMR – basal metabolic rate, Soma – somatic or vegetative growth, REB – reproductive energy budget.

To maintain energy reserves, individuals consume nutrients based on the photosynthetic rate, which is calculated using the relationship between plant mass and photosynthetic rate from the model of Hu et al. (2021). However, if the available nutrients in the landscape are limited, individuals consume what is available. Subsequently, individuals allocate part of their acquired energy and weight to basal metabolic maintenance. If individuals cannot mobilize enough energy to cover this maintenance cost, they die. The remaining energy uptake is divided into somatic growth and reproductive energy budget based on the Dynamic Energy Budget (DEB) models. The proportion of assimilated energy allocated to the soma is denoted as κ, while 1 − κ represents the proportion allocated to maturity maintenance, development, or reproduction (Kooijman 2009, Martin et al., 2012). Thus, the assimilated energy reserve from photosynthesis is allocated to three compartments: basal metabolic maintenance, somatic growth, and reproductive energy budget growth. We refer to this allocation ratio as metabolic efficiency. By altering the metabolic efficiency for tetraploids, we examine the effects of different metabolic efficiencies on population dynamics and tetraploid establishment. If an individual does not acquire enough energy for basal metabolism during a day, it experiences weight loss.

Energy for reproduction is accumulated over 100 days, representing a growing season. At the end of this period, all individuals with non-empty reproductive energy budget (REB) attempt to produce offspring – seeds. Seeds are initialized with seed size sampled from a normal distribution. During seed production, the reproducing individual’s REB decreases by the size of the seed being produced. As the modeled species is semelparous, individuals die after reproduction. In simulations of annual plants, individuals that fail to reproduce due to lack of REB also die.

During reproduction of diploids, each seed has a chance of undergoing polyploid formation. Instead of assuming a simple distribution with mean, we chose the polyploid formation rate to be Beta distributed parameter with mean 4.76%, shape parameters α = 2 and β = 40, based on previous findings (Ramsey 2007, Kreiner et al. 2017a). This specific distribution and parameterization were chosen to reflect the observations that the distribution of naturally forming unreduced gametes rate is heavily zero-inflated and that asexual species have a higher mean than sexual or mixed-mating systems (Ramsey 2007, Kreiner et al. 2017a). For simplicity, when tetraploid individuals reproduce, we assume they can only produce tetraploid offspring.

### Initialization

Per parameter combination, 10 simulations were run. At the start of a simulation, seeds were introduced with an average density of 1 individual per cell (a total of 1600 seeds). The size of the initialized seed was sampled from a normal distribution with mean 0.5 and standard deviation 0.05. The run time of all simulations was 10000 days (100 seasons). For exact initialization and parametrization of the model and tested scenarios, see the Supplementary Material.

### Consumer events

Here we give an overview of the events happening in the model and most important equations relating metabolism and size of the individuals. In Supplementary Material, we explain in detail.

#### Nutrient consumption and photosynthesis

The amount of nutrients taken up per day for an individual is determined as in equation 1:

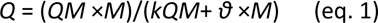

Where *Q* is the photosynthetic rate (g/day) and *M* is the plant mass (g). *QM, kQM* and *θ* are parameters stemming from the model of Hu et al. (2021) which reflect the joint effect of mass and limiting nutrients on photosynthesis. Based on their results for non-woody plants, the photosynthetic rate is calculated as

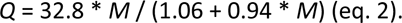

As the nutrients are limited (see section Resources), individuals at the same site experience nutrient competition to acquire these nutrients. Individuals experience scramble competition for nutrients. In compliance with metabolic theory, the photosynthetic rate, as the main metabolic rate considered here, is not dependent on the nutrient availability.

#### Metabolism and metabolic efficiency

In the model, the metabolic efficiency is defined as the ratio of energy allocated to basal metabolic maintenance, somatic growth, and reproductive energy budget growth. Initially, the diploid energy and nutrient usage is considered the most metabolically efficient, with a ratio of 36:54:10 (basal metabolic maintenance : somatic growth : reproductive energy budget). This ratio was chosen based on the input from DEB theory parameters, where in many species the proportion of assimilated energy allocated to the soma (κ) is around 0.8-0.9, while the proportion allocated to maturity maintenance and reproduction (1 – κ) is then 0.1-0.2. We choose the reproductive budget to be 0.1 or 10%, and then arbitrarily choose 36% for basal metabolic maintenance and 54% for somatic growth. To investigate the effects of different metabolic efficiencies on population dynamics and tetraploid establishment, the model explores scenarios where the metabolic efficiency of tetraploids deviates from that of diploids. Specifically, the model considers situations where the metabolic efficiency of polyploids is lower or higher than that of diploids. In the case of higher metabolic efficiency for polyploids, the model skews the ratio such that basal metabolic maintenance is lower than 36% of the total energy budget, allowing a greater proportion of energy to be invested in somatic growth and reproductive energy budget growth. This implies that polyploids allocate a larger portion of assimilated energy to growth and reproduction compared to diploids, and an expected (and trivial) result is the invasion of polyploids. Conversely, in the case of lower metabolic efficiency for polyploids, the model skews the ratio such that basal metabolic maintenance is higher than 36% of the total energy budget, potentially at the expense of reproductive energy budget growth. This means that a larger proportion of the assimilated energy is allocated to basal metabolic maintenance in polyploids, resulting in a reduced proportion available for somatic growth and reproductive energy budget growth. We explore the consequences of metabolic deficiency in polyploids on the dynamics, i.e. whether this reduction still allows polyploids to survive and establish in the ancestral diploid population in this model’s settings. Tested reduction of metabolic efficiency in polyploids range to −10% in comparison to the fixed diploid metabolic efficiency.

The model incorporates the idea that polyploids may exhibit immediate phenotypic differentiation, as observed in some species like *Heuchera grossulariifolia* (Oswald & Nuismer, 2011). This differentiation could result in different metabolic efficiencies for polyploids compared to diploids immediately upon formation. By exploring different metabolic efficiency scenarios, the model aims to capture potential variations in the metabolic strategies of polyploids and their implications for population dynamics and the establishment of tetraploids.

#### Reproduction

Energy for reproduction is collected during 100 days which comprise a growing season, after which all individuals with non-empty reproductive energy budget (REB) try to produce offspring – seeds. Seeds are initialized with seed size sampled from a normal distribution (mean 0.5, st dev 0.05). During seed production, the reproducing individual loses an amount of energy and size equivalent to the size of the seed from their REB. As each seed is initialized during reproduction of diploids, there is a chance of undergoing polyploid formation. For simplicity, when tetraploid individuals reproduce, we assume they can only produce tetraploid offspring, and seed sizes are assumed to be same as for diploid seeds.

Seed dispersal is also incorporated in the model and there is no cost of dispersal. Dispersal is random to the adjacent cells (nearest-neighbor dispersal). As we base this model on plants which are sessile organisms, there is no movement except this seed dispersal.

In addition to testing the impact of metabolic efficiency changes in polyploids, we also investigate the impact of different reproductive strategies. Perenniality is assumed to be beneficial for emerging polyploids, as this strategy could reduce MCE by providing a longer time-window for reproduction. It is however unclear whether perenniality is beneficial for polyploids in this model where the dynamics depends on metabolic efficiencies and competition for nutrients. In that regards we simulate scenarios:

- Case 1 - all individuals are annuals so whether or not they have filled reproductive energy budget after 100 days, they die.
- Case 2: all individuals are perennials, which means that they are allowed to live through multiple seasons until they can produce offspring, unless they die in the meantime from starvation or background mortality.
- Case 3: polyploids are perennial and diploids are annual, meaning polyploids can live for multiple seasons and have multiple opportunities for reproduction, whereas diploids die after a single growing season.

In total, simulations are done for metabolic efficiency where polyploids can be as efficient as diploids or less or more, for each of the three cases of annual/perennial reproductive strategies.

### Polyploid formation

To determine the rate of polyploid formation, the model incorporates empirical estimates of unreduced gamete formation. Studies in Brassicaceae have shown that unreduced gamete production varies among individuals, with an average rate of 2.5% and high individual-level variation (Kreiner et al. 2017a). The distribution of unreduced gamete formation rates is heavily zero-inflated (Ramsey 2007, Bretagnolle & Thompson 1995), and asexual species tend to have a higher mean than sexual or mixed-mating systems (Kreiner et al. 2017a, Kreiner et al. 2017b). To reflect these findings, the model uses a Beta distribution parameter with a mean of 4.76% and shape parameters *α* = 2 and *β* = 40. This distribution captures the zero-inflated nature of polyploid formation rates and the higher mean observed in asexual species.

### Data analysis

During each simulation, we tracked changes in the number of individuals (total, diploids, polyploids), amount of nutrients per cell, sizes of individuals and their REB. We also tracked whether the polyploid individuals are from diploid parents or from previously formed polyploids successfully reproducing. The results presented are based on 10 independent runs for each parameter set.

### Sensitivity analysis

Sensitivity analysis was conducted, and details on the tested parameters are available in the Supplementary Material 1.

## Results

### Polyploid invasibility and population dynamics

We undertake a comprehensive investigation into the potential for polyploid establishment and invasion by examining the final proportion of tetraploid individuals across a spectrum of metabolic efficiencies, as compared to diploids (Figure 2. This analysis shows that tetraploids can achieve dominance within the population even with metabolic deficiencies up to 5% (right side of Figure 2). The scenarios of annuals and perennials show minimal differences (Figure 2). However, when tetraploids are perennials opposing annual diploids (purple in Figure 2), invasion of tetraploids becomes feasible across a larger portion of the parameter space. Overall, the parameter space for polyploid-diploid coexistence is narrow. A closer look at the population dynamics over time (Figure 3) shows that in scenarios with failed polyploid establishment (metabolic costs of 10% as example), population sizes approach those as if polyploidy would not occur (Figure S1, S2).

**Figure 2.**
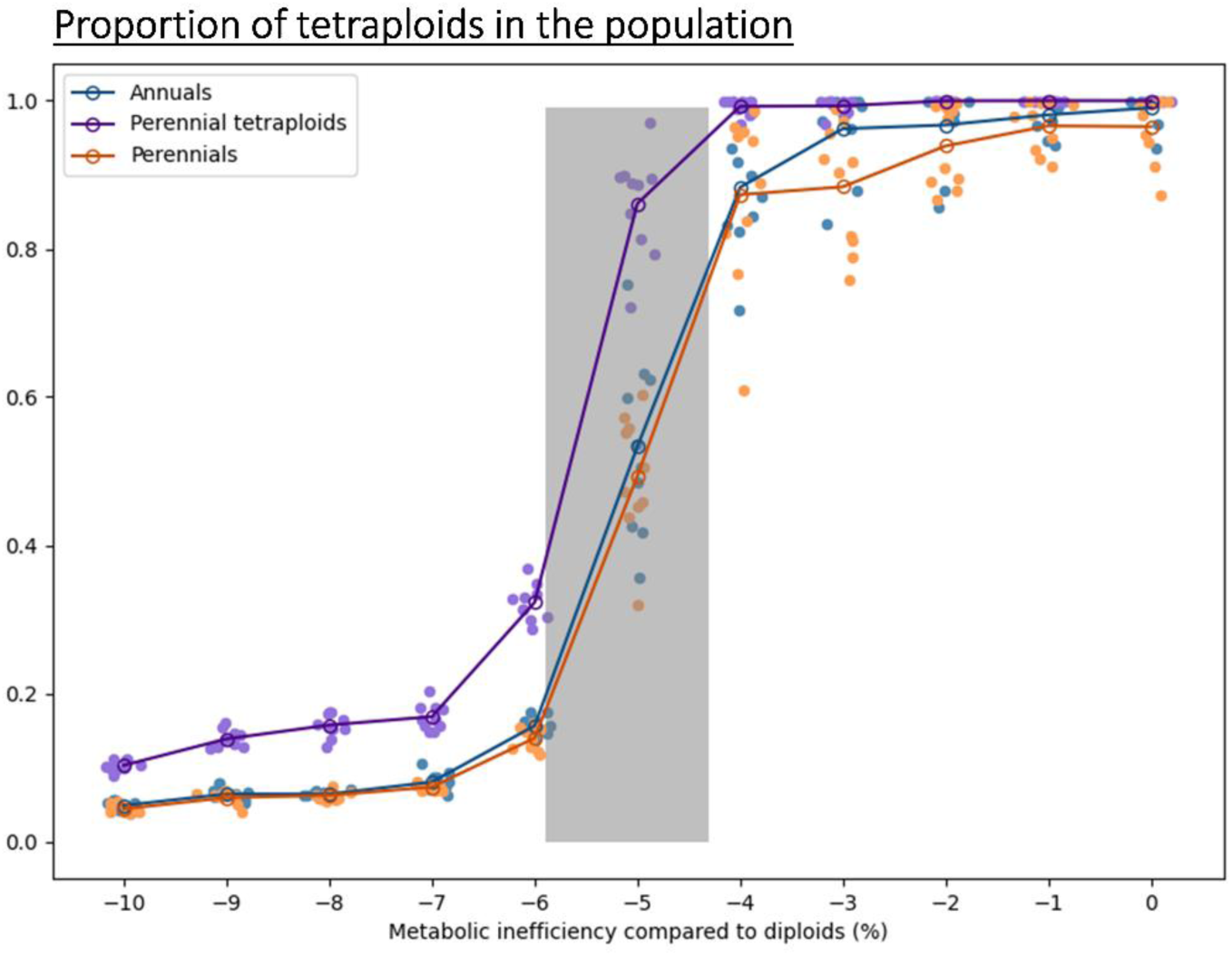
Proportion of tetraploid individuals across metabolic inefficiencies of tetraploids in comparison to diploids. Results are shown for the three simulated scenarios: all individuals annual in blue, all individuals perennial in orange and tetraploid perennials in purple. Independent simulation runs are represented by scattered dots, and darker shades are the means. Notice the grey shaded area on the plot which represents the zone where coexistence is observed.

**Figure 3.**
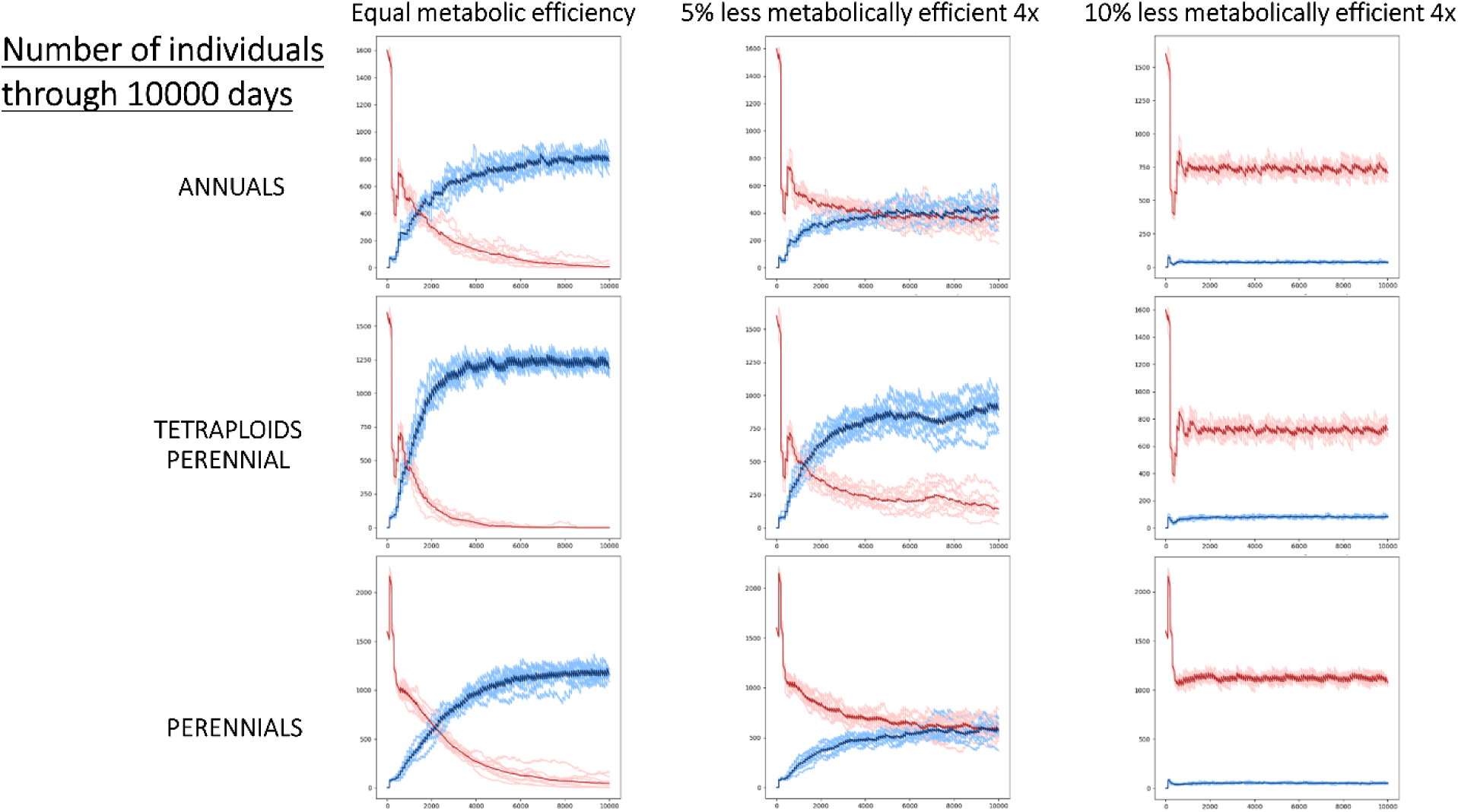
Number of individuals through 10,000 days, across simulated scenarios and metabolic efficiencies. Color of lines are differentiating between diploids (depicted in red) and tetraploids (depicted in blue). Different rows correspond to the 3 life history scenarios, with the top row corresponding to the annuals scenario, the bottom row to the perennials scenario, and the middle to the scenario where only tetraploids exhibit perenniality. Different columns represent tested metabolic efficiencies, with the first column showing results when tetraploid have an equal metabolic efficiency as diploids, and the second and third columns are showing results when tetraploids are 5 and 10% less efficient than diploids, respectively. Notice that these plots illustrate the mean of independent simulation in darker color shades of red and blue, while the lighter shades are all the independent simulations.

Overall, tetraploid establishment and success is only possible when majority of tetraploid seeds are originating from tetraploid parents (Figure S3). Insights are not sensitive to the used probability distributions of emergence (Figure S4), or the implemented polyploid formation rate (Figure S5). When polyploid formation rate is reduced to half, i.e., with a Beta distribution mean of 2.38% instead of 4.76%, the results maintain (see Figure S5b), although the zone of coexistence becomes broader and shifts more to the right. This again shows that the invasion success of tetraploids increases with lower metabolic inefficiencies. The pivotal role of recurrent polyploid formation is substantiated by exploring scenarios with intermittent bursts of polyploid formation, in contrast to the constant polyploid formation (see Figure S6). This examination reveals that single-season polyploid formation hinders establishment, and even with ten seasons of polyploid formation, successful establishment remains challenging unless polyploids exhibit high metabolic efficiency.

These insights hold for both the annual, mixed and perennial scenario despite changes in overall reached carrying capacities. Under perenniality, the accumulation of individuals persisting for multiple seasons increases the carrying capacity (measured as number of individuals) with about 50% (from ∼800 to 1200; Figure 3, compare top and bottom row). Contrary to initial expectations, perenniality does not inherently confer benefits. In fact, as elucidated in Figure 3 (bottom left), perenniality often postpones the invasion of tetraploids. This phenomenon arises from the expanded pool of individuals attempting reproduction across multiple seasons, subsequently intensifying competition for nutrients among individuals.

### Size and age at maturity

While population dynamics eventually determine chances of polyploid invasion, size and age distributions may be of high ecological relevance and even driving these population dynamics. Weight distributions show a multimodal distribution (Fig. 4A), which can be attributed to distinct phases in individual development. Young individuals, such as seeds and seedlings, exhibit low weights. As the simulation progresses in time, competitively superior individuals thrive by outcompeting others for nutrients, leading to substantially higher weights. Weight distributions align closely when metabolic efficiency is similar for diploids and tetraplods. Conversely, when the metabolic efficiency of tetraploids decreases, the weights of both reproducing and non-reproducing tetraploid individuals diminishes significantly. Noteworthy variations, however, manifest between the annual and perennial life history scenarios. When all individuals are perennial (third column of Figure 4A), an additional prominent peak emerges in the high weight distribution part. This peak stems from perennial individuals that have grown and aged over multiple years, benefiting from the capacity to persist even without reproduction. This peak is also prominent when tetraploids are allowed to adopt perenniality (second column of Figure 4). The high weight peak disappears when tetraploids are perennial and 10% less metabolically efficient than diploids. This highlights the challenges faced by tetraploids with diminished metabolic efficiency, as their reproduction success remains minimal, preventing their survival over multiple years under such inefficiency. Intriguingly, these distinctive peaks disappear when only reproducing individuals are considered (Figure 4B), hence demonstrating the persistent presence of non-reproducing individuals in the population.

**Figure 4.**
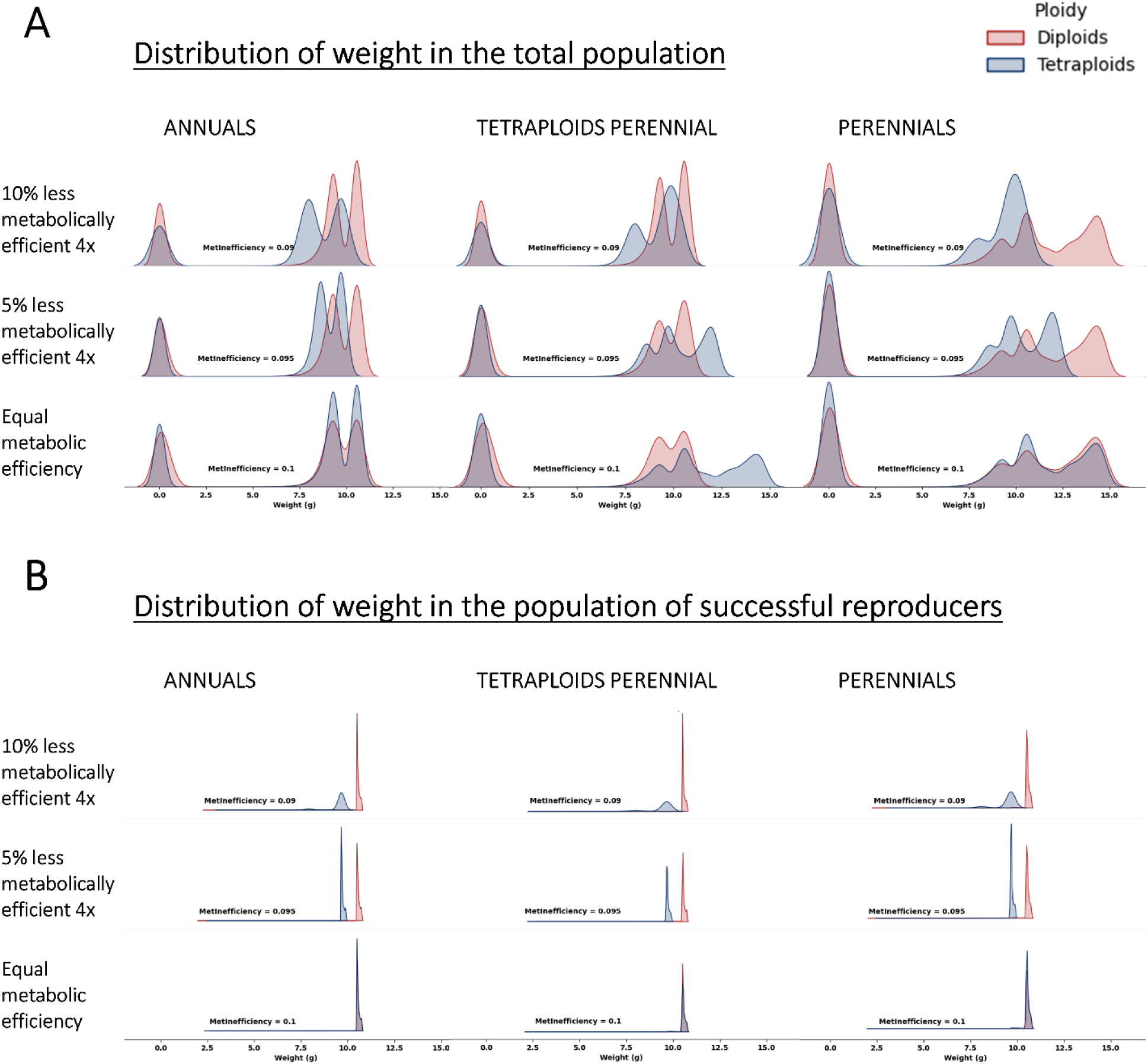
Distribution of individual weight across metabolic efficiencies and life history scenarios. A) shows the weights of all individuals, whereas B) shows only the weights of individuals which reached maturity and successfully reproduced. Different rows correspond to the tested metabolic inefficiencies of tetraploids in comparison to diploids, with the top and middle rows where tetraploids are 10% and 5% less efficient, respectively, and the bottom row where they are equal. Different columns correspond to the tested life history scenarios, with the first column corresponding to the annuals scenario, the second to the scenario where only tetraploids exhibit perenniality, and the third to the perennials scenario. Notice that red colors represent diploids and blue represent tetraploids.

In our simulations, annuals adhere to a single-year life cycle, relegating them to a single season for both living and reproduction. Consequently, the “all annuals” scenario is absent from the result of this section (Figure 5). Perennials are granted the potential to persist over multiple years until achieving successful reproduction or succumbing to chance or scarcity-induced death. The average ages of individuals typically exceed four seasons (Fig 5A). Notably, while the figure presents average ages, the simulated populations do encompass significantly older individuals, as highlighted in Figure S7. Very low metabolic efficiency in tetraploids extends the average time required for reproduction (Figure 5B), but on average, populations consist of old individuals that die-off without reproducing. In cases where only tetraploids adopt perenniality (even in the presence of tetraploid metabolic inefficiencies; see higher), age to reproduction is low and on average close to 1. This phenomenon arises from intra-cytotype competition dynamics, where individuals displaying swift growth and efficient utilization of the reproductive energy budget dominate the pool of reproducers. In line with the analysis of size dynamics (Fig 4), we show that individuals that persist for multiple seasons grow taller but do achieve successful reproduction.

**Figure 5.**
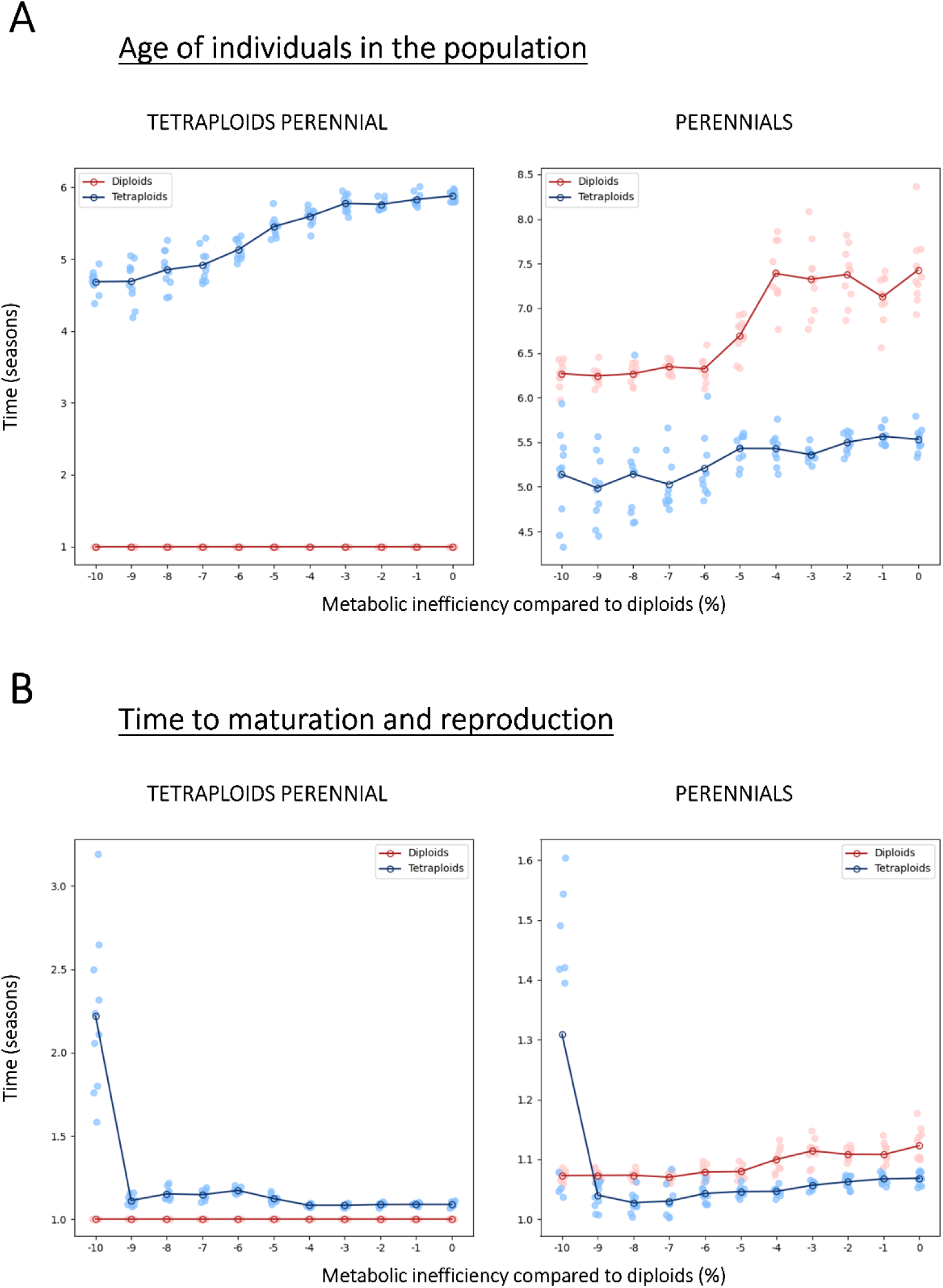
Age structure of diploid and tetraploid individuals across simulated metabolic inefficiencies of tetraploids and life history strategies. A) shows the average ages of all individuals, whereas B) shows average ages of only the individuals which matured and successfully reproduced. Color of lines are differentiating between diploids (depicted in red) and tetraploids (depicted in blue). Different plots correspond to the life history scenarios, with the left one corresponding to the scenario where only tetraploids exhibit perenniality and the right to the perennials scenario. Notice that these plots illustrate the mean of multiple simulations in darker color shades of red and blue, while the lighter shades are the independent simulations, and that annuals scenarios is not shown as there all the ages are 1 season.

A similar trend emerges in scenarios where all individuals are capable of exhibiting perennial traits. Here, intra-cytotype competition further accelerates the time to reproduction, aligning it more closely with the rapidity characteristic of annuals. The interplay between diploids and tetraploids introduces an additional layer of competition (inter-cytotype competition), encouraging tetraploids to prioritize faster maturation, especially when confronted with metabolic inefficiency. Essentially, the need to compensate for inefficiency propels successful individuals towards swift reproduction.

## Discussion

To better understand the eco-evolutionary dynamics of mixed ploidy populations, we used individual-based modeling (IBM), focused on the establishment from the perspective of invasion and intercytotype competition. IBM has been proven useful for studying emergent behavior and adaptation in changing environment, as it allows for the inclusion of a wide range of processes and interactions in a model. The model defines the processes and states in the system, while trait data of real biological systems can be used to define parameters and threshold states (Kearney et al. 2021). Although several models have been devised for studying polyploidy and its consequences (Griswold 2021, Fowler & Levin 2016, Suda & Herben 2013, Rodriguez 1996, Van Dijk & Bijlsma 1994, Felber 1991, Fowler & Levin 1984, Levin 1975), we devise a new model of polyploid emergence to study the effects of size and metabolic differences between ploidies on the population dynamics.

Population dynamics can be derived from relations of size and metabolism (metabolic theory of ecology, Price et al. 2010), and from intake and use of energy and resources (Dynamic Energy Budget (DEB) theory, Kooijman 2010). These general ecological theories have been used to make predictions about various biological phenomena on population and ecosystem levels, and here we employ them as a general framework for building a model about population where polyploidization happens and newly formed polyploids may or may not have metabolic consequences. We show that polyploid establishment and invasion is possible even with lower metabolic efficiency of the neopolyploids, but that it is highly conditional on the recurrence of polyploid formation. Contrary to prevailing assumptions, we demonstrated that perenniality does not inherently confer advantages for neopolyploids, but perenniality greatly changes the population structure and thereby impose increased intra- and intercytotype competition for nutrients. This competition forms the foundational basis driving subsequent population dynamics, influencing the potential success of polyploids. Our study’s unique contribution lies in being the first to explore the eco-evolutionary dynamics of polyploid establishment through the lens of basic ecological mechanisms.

We here deliberately focused on feedbacks from size, metabolic changes to life history as a sufficient criterion for the outcome of polyploid-diploid coexistence. Neopolyploids with metabolic efficiencies larger, equal or only slightly less than their diploid ancestors can successfully establish and invade the population. Metabolic inefficiency of more than 5% hampers polyploid establishment, emphasizing the need for real plant polyploids to adapt and counterbalance the costs associated with larger cell size and reduced metabolism. Such a metabolic efficiency arises in plants from a multitude of downstream cellular processes when genome doubling leads to genomic instability, mitotic and meiotic abnormalities, gene expression and epigenetic changes (Comai 2005). In plants, CO_2_ exchange rates are a well-known measure for basal metabolic processes. Such CO_2_ exchange rate declines as ploidal level increases in many species, and tetraploids average about 11% less than diploids (Levin 1983). In artificial colchicine-induced autopolyploids of *Arabidopsis thaliana*, differences in the concentrations of metabolites related to the tricarboxylic acid (TCA) cycle and the γ-aminobutyric acid (GABA) were found (Vergara et al. 2016), which could have important adaptive consequences for the ecology of diploids and polyploids due to the diverse functional roles of TCA and GABA metabolites. We here show that such metabolic decreases should not induce a direct paradox with respect to the polyploid persistence. Evidently, we expect species with polyploids having higher metabolic efficiencies, (e.g., *Beta vulgaris* (Levin 1983), or reported that CO_2_ fixation rates were higher in autotetraploid than in the diploid and in the naturally occurring polyploid series of *Festuca arundinacea* (Byrne et al. 1981)) to eventually outcompete their diploid ancestor. Conversely, any coexistence with diploids likely points at stabilizing processes of niche divergence, or equalizing processes that reduce any fitness advantage of the polyploid. In this respect, chlorophyll fluorescence parameters, an overall indicator of photosynthesis, were found to be higher in neotetraploids of *Jasione maritima* var. *maritima* (Siopa et al. (2020). However, this higher efficiency was not reflected in increased carbohydrates (total soluble sugars) and biomass production. Likely, neotetraploids invest more resources in defense mechanisms (e.g., increase of antioxidant battery) compromising growth, while diploids allocate most of their resources in growth. Likewise, in Castro et al. (2023) *Jasione maritima* diploids seemed to invest more into biomass and less in starch accumulation in comparison to tetraploids. The lack of such gigas effect were here explained by developmental tradeoffs.

The role of recurrent polyploid formation emerges as a significant driver of establishment success. Our model results also suggest that intermittent polyploid formation in not permissive for successful establishment, except in scenarios which require high metabolic efficiencies in the neopolyploids. Recurrent polyploidy is found in most studied plant systems (Soltis & Soltis 1999, Kolář et al. 2012, Shimizu-Inatsugi et al. 2009, Čertner et al. 2017), which is reflected in this model by having a chance to produce polyploid offspring in each reproductive trial. However, clear empirical data about the variation of unreduced gamete formation are lacking, which is why we choose Beta probability distribution with mean of about 4.76% to reflect insights from Ramsey 2007 and Kreiner et al. 2017.

Clonality and perenniality have been often found in polyploid taxa, but perenniality seems to be a weak contributor to polyploid evolution, in some Asterids clade at least (Van Drunen & Husband 2019). Diploid and autotetraploid plants have different growth rhythms, affecting their longevity and habit (Levin 1983). For example, an autotetraploid strain of *Zea mays* is perennial, while diploid maize is annual. Similar patterns are observed in species like *Eragrostis* and *Nasturtium*. Greenhouse trials with *Oryza punctata* showed that tetraploids survived 2-9 years, while diploids rarely lived beyond 1 year (Sano 1980). Greater longevity in tetraploids is also observed in *Trifolium* species (Frame et al. 1976) and *Medicago* (Small 2011). Due to associations of perenniality with polyploidy, and its theoretical benefit for reproductive assurance, we investigated scenarios with annuality vs. perenniality in our model. Contrary to previous assumptions about perenniality providing sufficient time and opportunity for newly formed polyploids (Rice et al. 2019), our study challenges the notion that perenniality inherently confers benefits to polyploid establishment as a reproductive reassurance mechanism. When diploid and tetraploid individuals are both allowed to be perennial, invasion of tetraploids is feasible across same parameter space as when all the individuals are annual. Perenniality often delays the invasion of tetraploids, attributed to intensified competition for resources arising from the prolonged presence of individuals attempting reproduction over multiple seasons. Ecological factors, nutrient competition specifically, affected the model outcomes more than the life history strategies, in line with model of Fowler & Levin (2016). Being perennial does not seem to be beneficial when the environment (in this case nutrient availability) is predictable, because faster life histories like annuals outcompete in such conditions, when semelparity is considered (Paniw et al. 2018).

The exploration of size and age at maturity provides insight into the feedbacks between size-dependent photosynthesis, metabolic efficiencies, and reproduction. The observed multimodal weight distributions highlight distinct phases of individual development, but also that metabolically deficient tetraploids only succeed to grow to smaller weights. This pattern persists for size at maturation. Metabolic efficiency plays a pivotal role in shaping weight distributions, with decreasing efficiencies leading to diminished weights among both reproducing and non-reproducing tetraploid individuals. The influence of metabolic efficiency on reproduction success becomes evident in scenarios where tetraploids are perennial and 10% less metabolically efficient than diploids. Perennial individuals tend to grow to high weights but rarely reach maturity and reproductive capability in this model. This emphasizes the trade-offs and challenges faced by tetraploids with reduced metabolic efficiency, hindering their survival over multiple years. This is contrary to the expectation that larger individuals are more fit than smaller ones that grow slowly, have lower storage capacity and poorer recovery from phosphorus depletion (Malerba et al. 2018). Our findings contradict the typically observed larger sizes of polyploids in nature (Bomblies 2020). Other processes during development or advantages of larger sizes that overrule the metabolic costs (Malerba et al. 2018) and leading to niche differences are important drivers of size differences between polyploids and their ancestors. Only in some rare cases, for instance smaller fruits found in autopolyploids in a Cactaceae species (Cohen and Tel-Zur (2011), smaller sizes may thus be explained by metabolic processes.

The age structure analysis further reveals the importance of metabolic efficiency in determining the time to reproduction and confirms that perenniality is not beneficial, as due to the competition for nutrients, only fast, on average annual, individuals reproduce, while most individuals living for multiple seasons do grow in size, without contributing to reproduction (“living deads”). Neopolyploids with low metabolic efficiency exhibit extended time to reproduction, highlighting the implications of metabolic costs on reproductive timing. The interplay between intra- and intercytotype competition contributes to the dynamics of time to reproduction, with successful individuals prioritizing rapid maturation to compensate for inefficiency. So overall, we find that competition for nutrients is driving the population dynamics and influencing the success of tetraploids in this model. As smaller individuals require less nutrients, they are competitively superior in these competitive environments.

In conclusion, our study illuminates the complex interplay between metabolic efficiency, life history strategies, size-dependent dynamics, and reproduction success in polyploid establishment. The insights derived from our mechanistic modeling approach pave the way for a paradigm shift in polyploidy research, emphasizing the importance of integrating ecophysiological insights to elucidate long-standing questions about polyploid establishment (Soltis et al., 2016). Our study exemplifies the potential of mechanistic modeling to bridge ecological and evolutionary perspectives in understanding polyploid establishment. By integrating ecophysiological insights with ecological modeling, we offer a holistic view of the factors influencing polyploid success. This approach transcends traditional population genetics models, providing a more nuanced understanding of polyploidy’s ecological implications. In addition, parameters of the model presented here, like photosynthetic rate and size, are measurable for real systems so predictions made here can be tested in the future.

## Acknowledgements

YVdP acknowledges funding from the European Research Council (ERC) under the European Union’s Horizon 2020 research and innovation program (No. 833522) and from Ghent University (Methusalem funding, BOF.MET.2021.0005.01).

## Supplementary material 1

This model description follow the ODD protocol (Overview, Design concepts, Details) for describing individual-based models (Grimm et al. 2006, 2010, 2020).

### Purpose

Our aim was to investigate how polyploid establishment in a diploid population is influenced by differences in metabolic efficiencies and life history strategies. We accomplished this by employing a mechanistic, individual-based model (IBM) grounded in consumer-resource dynamics. We determined the parameter combinations that facilitate polyploid establishment, coexistence and invasion.

By applying an IBM, we successfully incorporated variations in polyploid offspring formation and stochasticity within our model. The individuals’ ploidy levels are categorized as diploid and tetrploid, without explicit representation of the underlying genome. This approach, coupled with the assumption of clonal reproduction, implies that our results can be interpreted both for asexually reproducing species and females of a sexual species which does not suffer from pollen limitation. Other interpretations are also viable, also depending on the ploidy levels considered (e.g. haploids with emerging diploids).

### Entities, state variable, and scales

#### The consumers

The consumer species is individually modeled and has the following state variables:

-ploidy: set to have values 2 or 4 to imitate polyploidization to tetraploids within a diploid species

-W0 (g): the seed mass of an individual

-W (g): referring to the current somatic mass of an individual

-ER (g): mass of the reproductive energy budget, which is how much of the body size can be invested in reproduction

-CM (g/day): consumption, mass of the nutrients that should be consumed during a day, calculated for the individual’s total body mass (W+ER), based on Hu et al. (2021) photosynthetic rate prediction for non-woody plants

-nutrient allocation ratio: the total mass of the consumed nutrients within a day (CM) is used for basal metabolic maintenance (BM), then somatic growth (SG) and then reproductive energy budget growth (REB). Based on Hu et al. (2021), allocation for diploid individuals is set to be 36 BM: 54 SG: 10 REB. For tetraploid individuals, this allocation ratio changes in order to simulate higher (lower BM and higher REB) or lower (higher BM and lower REB) metabolic efficiency.

-birthday and age (days): referring to the age, important for simulations of perennial life cycles as they might live for several seasons

-parent ploidy: important for tetraploid emergence estimation as tetraploids may have diploid or tetraploid parents

-x and y coordinate

#### The landscape

The landscape is a lattice of 40x40 cells, where each cell has certain amount of resources (nutrients), represented as mass in grams. The edges of the landscape are not wrapped so there can be effect on the seed dispersal. A cell is defined by following state variables:

-R0 (g): initial amount of the nutrients. Each cell is initialized with 1 g of resources, as an abstraction of main nutrients needed for plant growth like nitrogen and phosphorus.

-r(g/day): growth speed of the nutrients. Nutrients are growing linearly each day by 1 g (green shaded box in Figure S1).

-Rmax(g): maximum amount of nutrients, 10 g in all cells. Nutrients availability is then auto-correlated in time.

#### Scales

Seed dispersal is the sole form of movement considered in the model, drawing inspiration from sessile plant species. Given that the primary focus of this model was to examine a diploid population with emerging polyploids, the landscape size was intentionally kept relatively small. Seed dispersal was confined to the nearest neighbor cells. Each time step corresponds to one day, and a growth season of 100 days was used as the timeframe for reproduction opportunities. All simulations were conducted over 10,000 time steps, equivalent to 100 seasons.

### Process overview and scheduling

The utilized model is a spatially explicit, discrete-time framework featuring daily growth events and seasonal reproduction events. Each time step corresponds to a single day in the consumer’s lifecycle, and a season is represented by a span of 100 days. In line with a plant species-based approach, we adopt a semelparous reproduction strategy, wherein reproduction occurs once in an individual’s lifetime following the conclusion of a 100-day growth season. Nutrient availability is conceptualized as nitrogen- and phosphorus-limited carbon uptake for photosynthesis.

**Figure S1:**
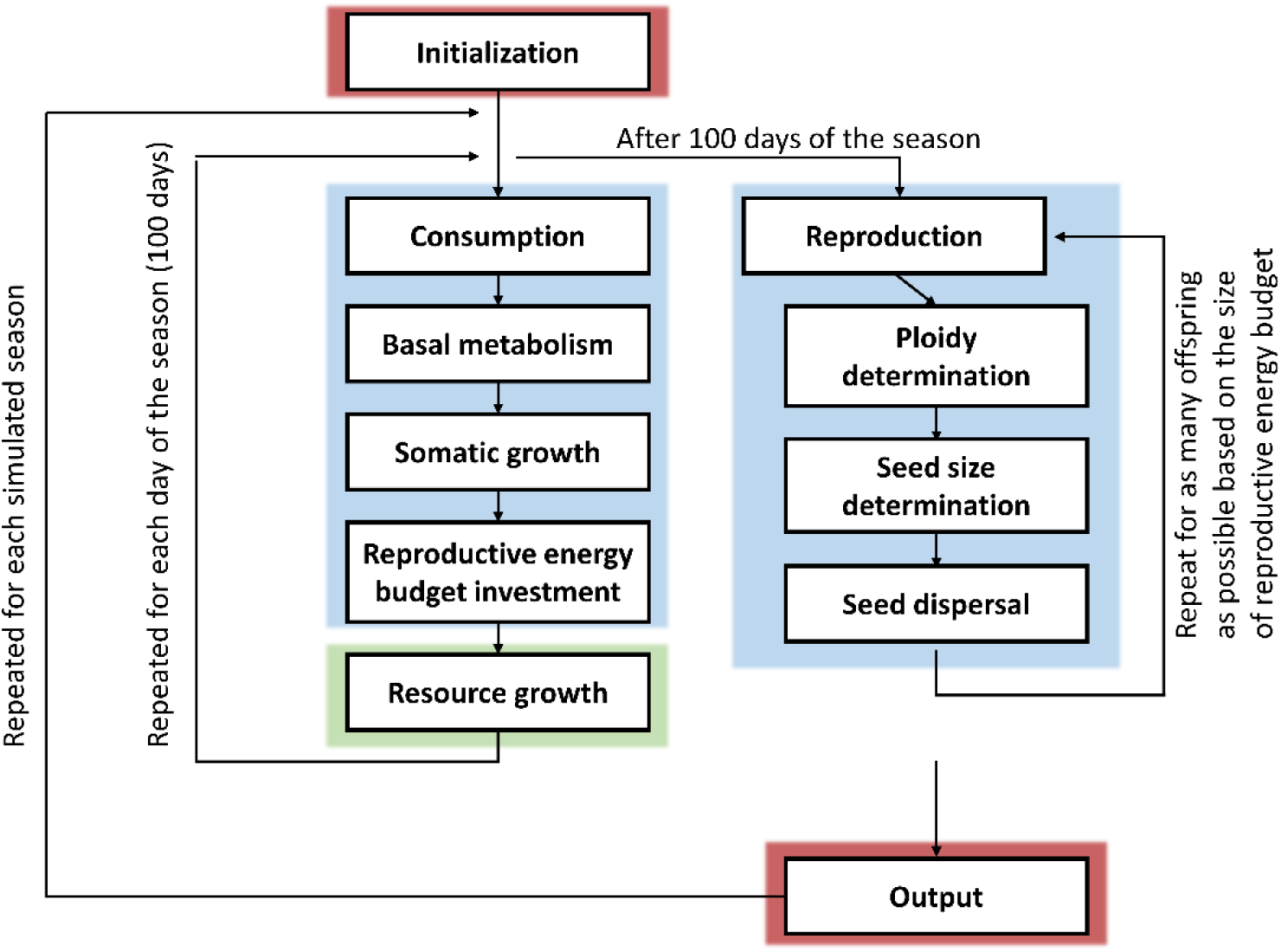
Depiction of all events within the model. Initialization and output generation are regulated at the level of the model (red boxes). Blue boxes represent events of the consumers and the green box represents the nutrients growth.

Individuals reproduce asexually, driven exclusively by their reproductive energy budget size that has accumulated over the course of the season. Movement is not allowed as plants are sessile organisms, but there is dispersal of the newly formed offspring (seed dispersal, randomly to one of the nearest neighboring cells or parental cell).

During each day, every individual attempts to consume nutrients from the landscape. The amount of nutrients that individual tries to consume is related to its current weight, based on Hu et al. 2021. Their model incorporates a relationship between photosynthetic rate and carbon uptake in relation to plant mass, accounting for nitrogen and phosphorus as limiting factors for plant growth. In instances of nutrient scarcity, individuals undergo a reduction in weight as they allocate energy to basal metabolic maintenance, essential for daily survival. Consequently, energy allocated to growth leads to reversible increases in body size. Consequently, individuals may perish due to “starvation.”

When an adequate nutrient supply is available, an individual’s sequence of actions involves deducting the basal metabolic maintenance “fee,” subsequently allocating nutrients to somatic growth, and ultimately contributing to the reproductive energy budget. At the close of each day, nutrient levels are restored.

At the end of the season (i.e. after 100 days), individuals reproduce if their reproductive energy budget is not depleted, following which they perish. Reproducing individual allocates the REB to producing seeds of certain seed sizes (sampled from normal distribution with mean 0.5 and st dev 0.05), continuing this allocation until the budget is exhausted. Diploid individuals have the potential to produce tetraploid offspring, with the probability of this occurrence being sampled from a Beta distribution with mean 4.76%, shape parameters α = 2 and β = 40. Conversely, tetraploid individuals are restricted to only producing tetraploid offspring.

The sequence in which individuals consume does not affect the final outcome, as their order is randomized before execution of the daily events.

### Design concepts

#### Basic principles

One of the most commonly observed phenotypic consequence of polyploidization is an increase in cell size and overall body size (Stebbins 1971, Bomblies 2020). Body size stands as a crucial attribute in the ecophysiology of organisms. The Metabolic Theory of Ecology established the correlation between body size and numerous functional traits (Peters, 1983; Brown et al. 2004, Reich et al. 2006, Price et al. 2010), such as basal metabolic rate, ingestion rate, and developmental time. A fundamental tenet in biology is that metabolic rates scaling allometrically to body size, often approximated by a scaling exponent of 0.75 (Kleiber 1932). Notably, in plant and phytoplankton taxa, respiratory rate scaling appears to deviate from the animal pattern, exhibiting a higher exponent approaching 1 (isometry, Reich et al. 2006; Lopez-Sandoval et al. 2014), even though other indicators of metabolism, such as biomass production rate, adhere to a body size exponent of 0.75 (Niklas & Enquist 2001). Given this variability, for the purposes of our model, we opted to employ a distinct metabolic framework proposed by Hu et al. (2021). This model captures the relationship between photosynthetic rate and body size while also accounting for nutrient considerations. The model’s fit yielded superior explanatory power for plant data compared to the allometric scaling model, prompting our decision to adopt it. We employ their estimation for non-woody plants as a formula governing the daily amount of photosynthesis (consumption) in a plant.

Due to having twice the amount of DNA as their diploid predecessors, tetraploids are postulated to necessitate increased energy and nutrient allocation for basal metabolic maintenance. However, this heightened allocation could potentially limit nutrients available for somatic and reproductive growth. As a reference, we regard diploid energy and nutrient utilization as the most metabolically efficient, characterized by a distribution of 36% for basal metabolic maintenance, 54% for somatic growth, and 10% for the reproductive energy budget (36:54:10). Consequently, we investigate the implications of varying metabolic efficiencies in tetraploids on population dynamics and the establishment of tetraploids. This is achieved by altering the 36:54:10 ratio to reflect lower metabolic efficiencies in tetraploids—for instance, 37:54:9, indicating a 10% reduction in efficiency compared to diploids.

The notion that polyploids may exhibit suppressed metabolism was postulated by Cavalier-Smith (1978), who drew from data obtained from *Ribes satigrum*, *Datura stramonium*, *Raphanus sativus*, *Galeopsis pubescens*, *Solanum nudiflorum*, *Vitis vinifera*, and *Phlox drummondii*. In these cases, CO2 exchange rates decreased with increases in ploidy levels, possibly due to alterations in the geometry of polyploid cells. We explore the implications of these differences in metabolic efficiency across three distinct life history strategies: annual individuals, perennial individuals, and annual diploids coexisting with perennial tetraploids. The outcomes of these diverse models are then subjected to comparison.

While it is generally anticipated that polyploids would exhibit diminished metabolic rates in contrast to diploids, there are instances that deviate from this trend (Levin 1983), indicating potential increases in metabolic rates among polyploids. We consequently simulated these variations in metabolic efficiency as well.

#### Emergence

Population dynamics emerge from the behavior of the individuals, where daily life cycle is represented by empirical rules describing metabolism. Successfully reproducing diploids can have diploid and tetraploid offspring, as there is a probability (Beta distribution with mean 4.76%, shape parameters α = 2 and β = 40) for each offspring to be tetraploid. With no inherent differentiation between diploids and tetraploids, the tetraploid accumulation is anticipated due to their role as a sink.

Nutrient distribution undergoes spatial alterations as individuals consume nutrients daily, amplifying the significance of competition. Smaller individuals require fewer nutrients, which may allow multiple individuals in a cell to sustain growth and investment in the reproductive energy budget. Conversely, larger individuals exhibit heightened resistance to starvation, as their greater mass provides a buffer against nutrient scarcity. Disparities in metabolic efficiency between diploids and tetraploids are poised to shape population dynamics, contributing to the emergence of distinct body sizes.

Furthermore, variations in life history strategies are anticipated to yield diverse population dynamics. In the context of annuals, which experience only a single season, those individuals that secure optimal nutrients and allocate them to the reproductive energy budget attain the highest absolute fitness. In contrast, the perennial strategy boasts advantages as reproduction is postponed to subsequent seasons, permitting individuals to accumulate more nutrients and potentially augment their reproductive energy budget.

As a result, the distribution of body sizes will materialize as an outcome of nutrient availability and metabolic efficiencies, intertwined with life strategies. Together, these factors culminate in determining population dynamics, including the potential coexistence of tetraploids with diploids or the invasion of tetraploids.

#### Adaptation

In this model, body size and ploidy stand as the principal defining characteristics of individuals. The rate of consumption depends on body size, resulting in dynamic changes attributable to the allocation among basal metabolic maintenance, somatic growth, and the reproductive energy budget.

Within the confines of a shared cell, individuals engage in competition for limited resources. This competition yields instances where some individuals consume less than the optimal amount of nutrients, leading to a loss in their reproductive energy budget and somatic weight. It’s important to note that adaptation is not a feature of this model, as it lacks an inheritance mechanism.

#### Stochasticity

All demographic parameters are treated as probabilities, or are drawn from probability distributions. This approach was adopted to include demographic noise and emphasize population-level phenomena, rather than individual behavior.

#### Observation

During each time step, we recorded the counts of individuals (both diploid and tetraploid) and nutrient levels across all cells, alongside the weights of individuals. Upon reproduction, the ages and weights of reproducing individuals were also documented. The spatial distribution of individuals was visualized to provide a comprehensive perspective.

### Details

#### Initialization

For each parameter combination, 10 simulations were run, each with 10 000 time steps or 100 seasons. At the start of each simulation, 1600 diploids were introduced as seeds and randomly distributed across the grid cells of the habitat. Every grid cell was initiated with 1 gram of nutrients. The initial number of individuals was selected to maintain an average density is 1 individual per cell.

#### Input data

Seed sizes for each individual are drawn from a normal distribution with mean 0.5 and standard deviation 0.05. Background mortality was kept in all submodels, with rate of 5 individuals per 1000.

#### Submodels

Case 1 - all individuals are annuals so whether or not they have filled reproductive energy budget after 100 days, they die.

Case 2: all individuals are perennials, which means that they are allowed to live through multiple seasons until they can produce offspring, unless they die in the meantime.

Case 3: polyploids are perennial and diploids are annual, meaning polyploids can live for multiple seasons and have multiple opportunities for reproduction, whereas diploids die after a single growing season.

**Table S1.**
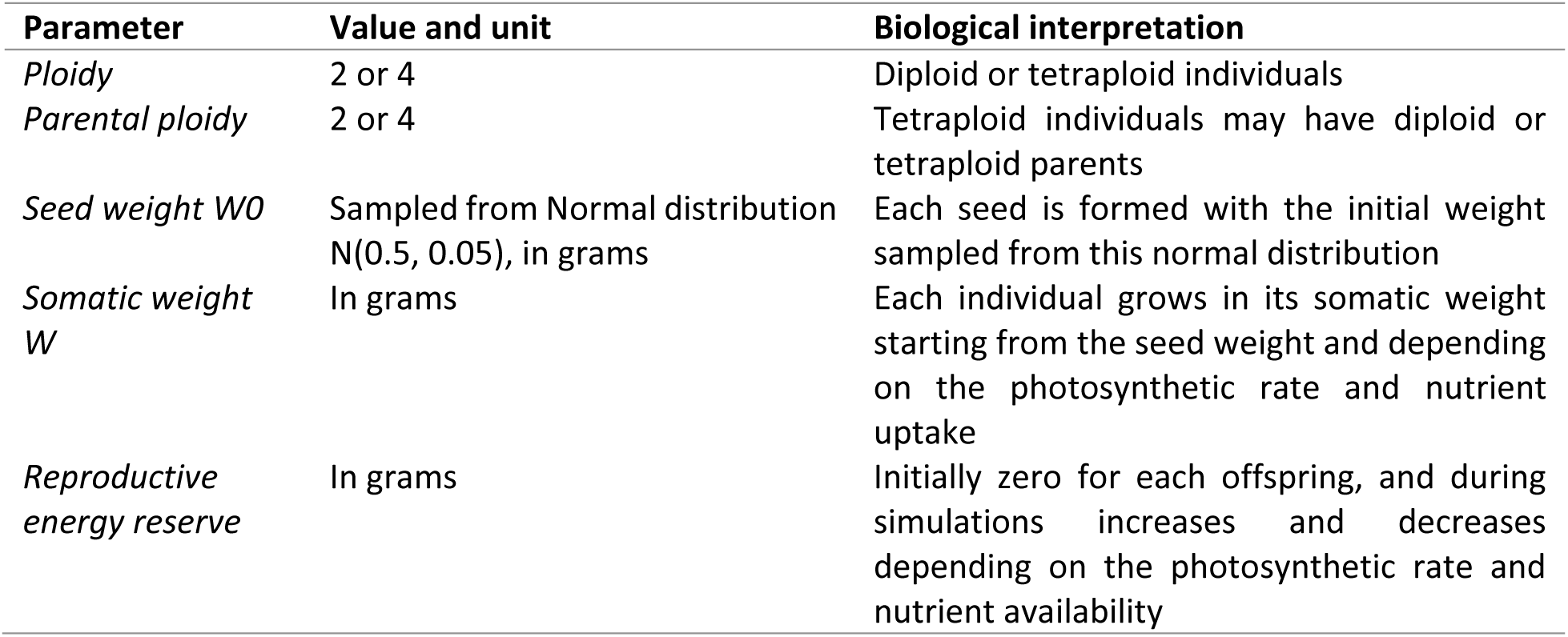
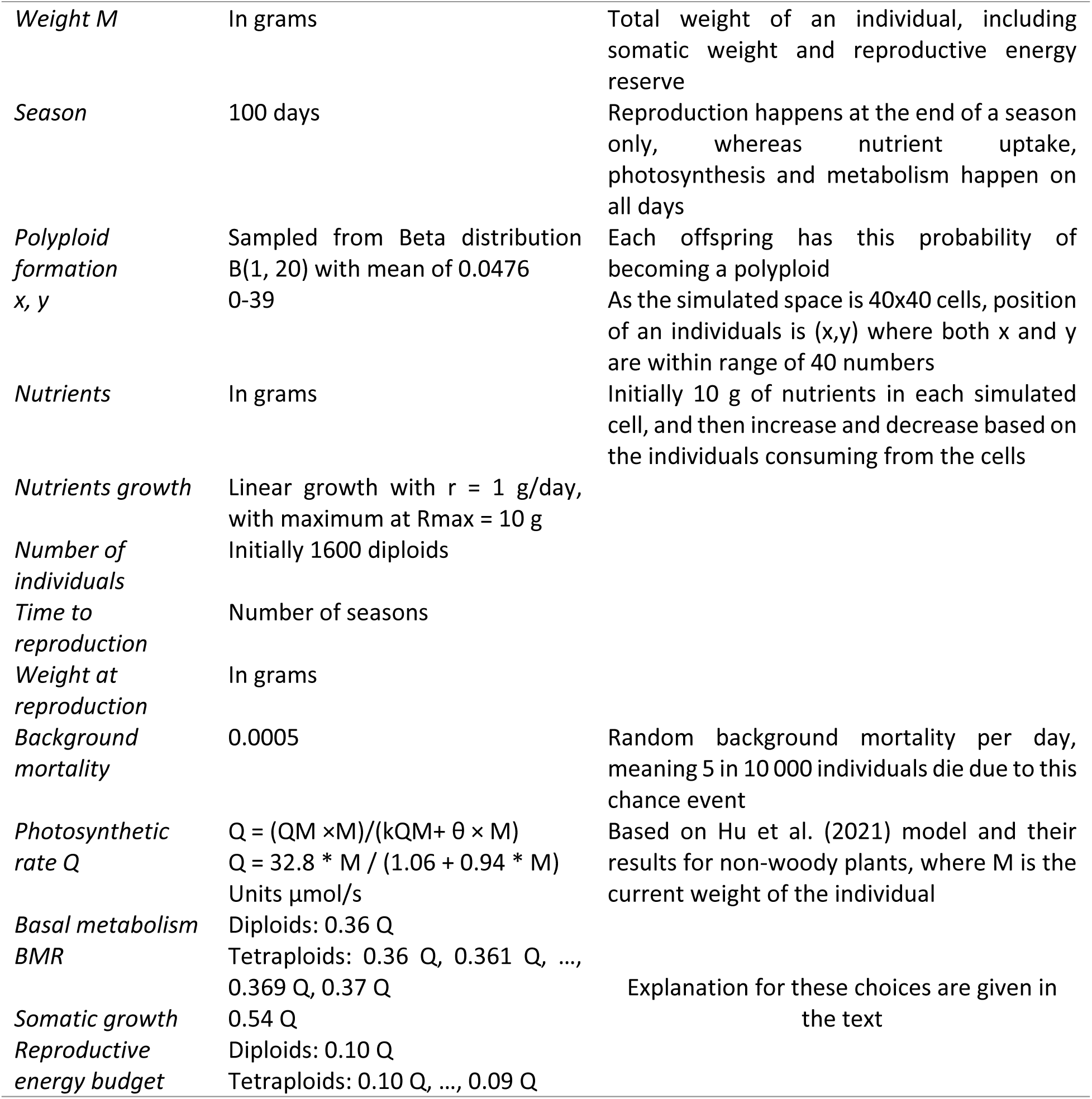
Set of parameters used in the model simulations developed in this study. The table contains the parameter name and symbol, the parameter values used, and a brief description of the biological interpretation.

## Supplementary material 2

**Figure S1.**
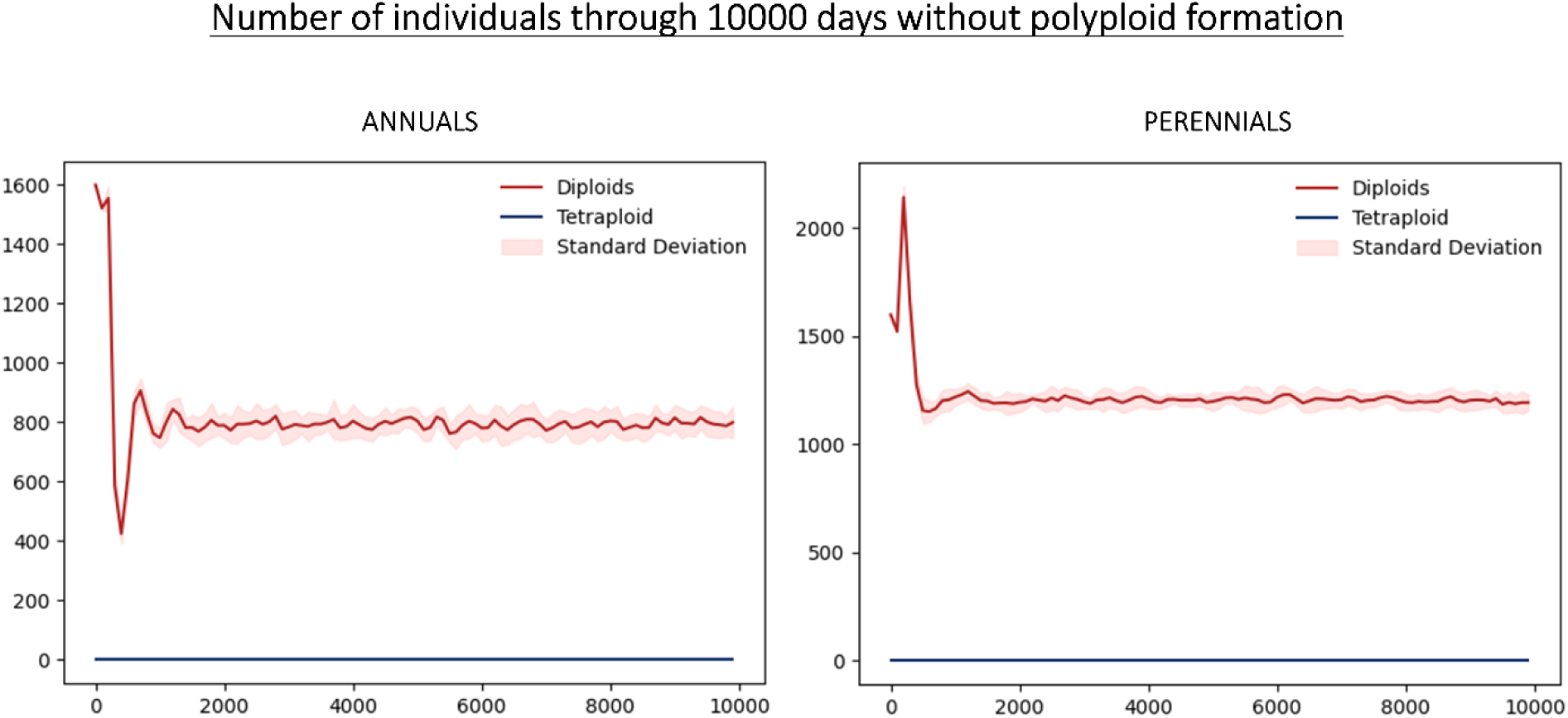
Number of individuals through 10,000 days, without polyploid formation allowed in the simulations. Color of lines are differentiating between diploids (depicted in red) and tetraploids (depicted in blue). Results are shown for the two opposing strategies, i.e. annuality and perenniality. Notice that these plots illustrate the mean of independent simulation in darker color shades, while the lighter shade is the standard deviation. Estimated carrying capacity after stabilization was 814 and 1237 for annuals and perennials, respectively.

**Figure S2.**
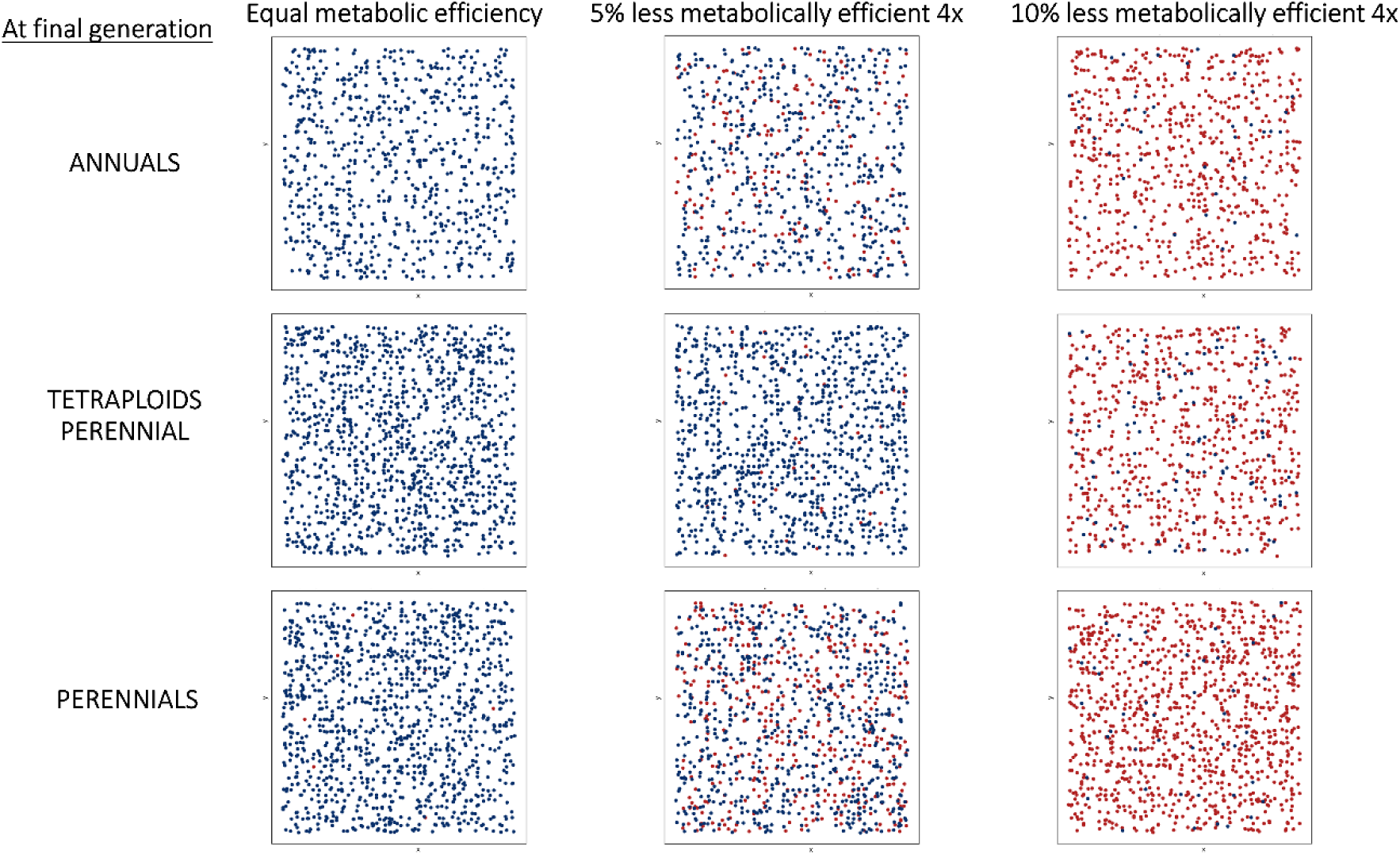
Spatial distribution of individuals across simulated scenarios and metabolic efficiencies. Individuals are represented as dots in x, y space, with color of dots differentiating between diploids (depicted as red dots) and tetraploids (depicted as blue dots). Different rows correspond to the 3 life history scenarios, with the top row corresponding to the annuals scenario, the bottom row to the perennials scenario, and the middle to the scenario where only tetraploids exhibit perenniality. Different columns represent tested metabolic efficiencies, with the first column showing results when tetraploid have an equal metabolic efficiency as diploids, and the second and third columns are showing results when tetraploids are 5 and 10% less efficient than diploids, respectively. Notice that these plots illustrate the results from a single run of the simulation, at the final generation (generation 100).

**Figure S3.**
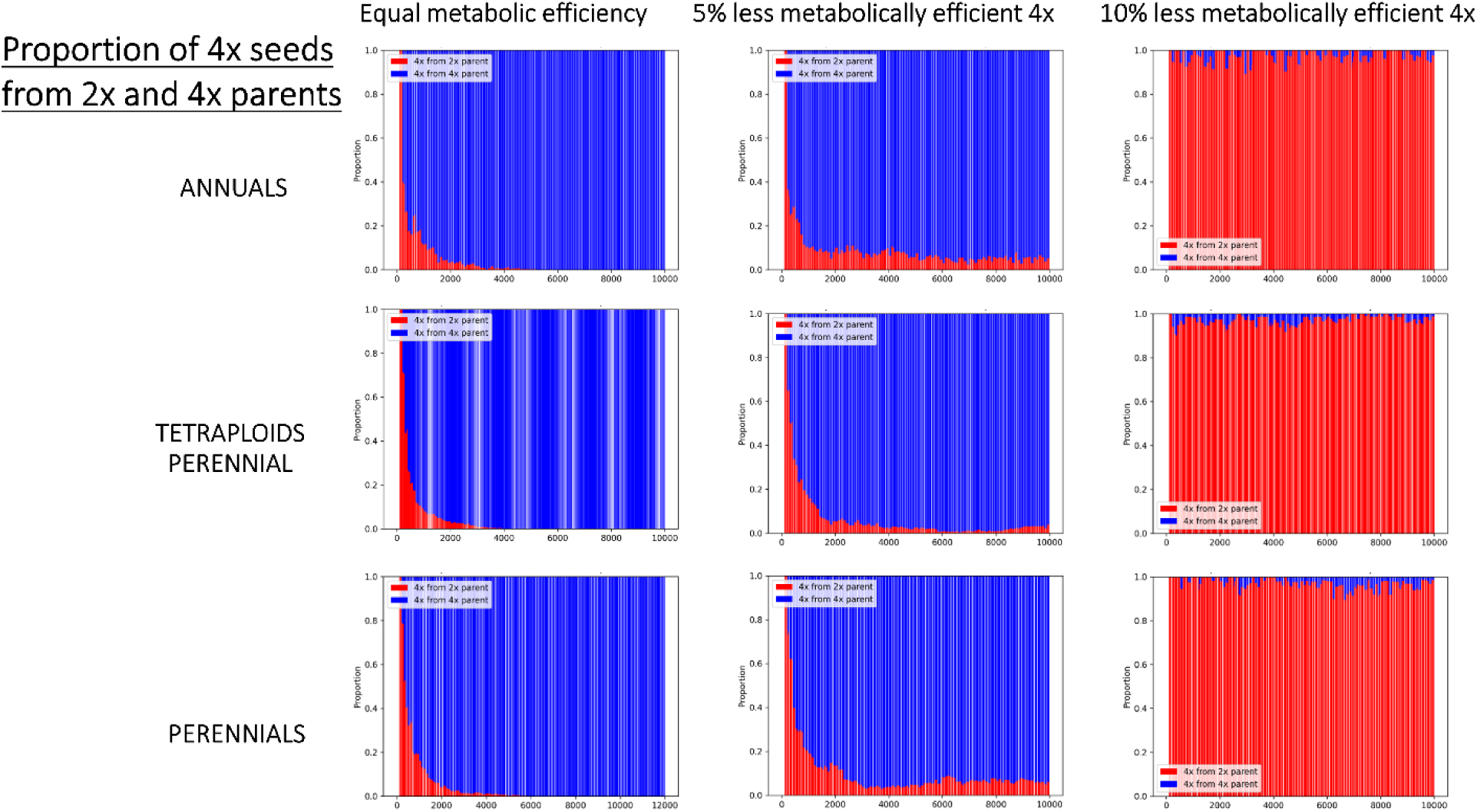
Proportion of tetraploid seeds from diploid and tetraploid parents through 10 000 days across simulated scenarios and metabolic efficiencies. Color of the bars on the histograms are differentiating between diploid (depicted in red) and tetraploid parents (depicted in blue). Different rows correspond to the 3 life history scenarios, with the top row corresponding to the annuals scenario, the bottom row to the perennials scenario, and the middle to the scenario where only tetraploids exhibit perenniality. Different columns represent tested metabolic efficiencies, with the first column showing results when tetraploid have an equal metabolic efficiency as diploids, and the second and third columns are showing results when tetraploids are 5 and 10% less efficient than diploids, respectively. Notice that these plots illustrate the results from a single run of the simulation.

**Figure S4.**
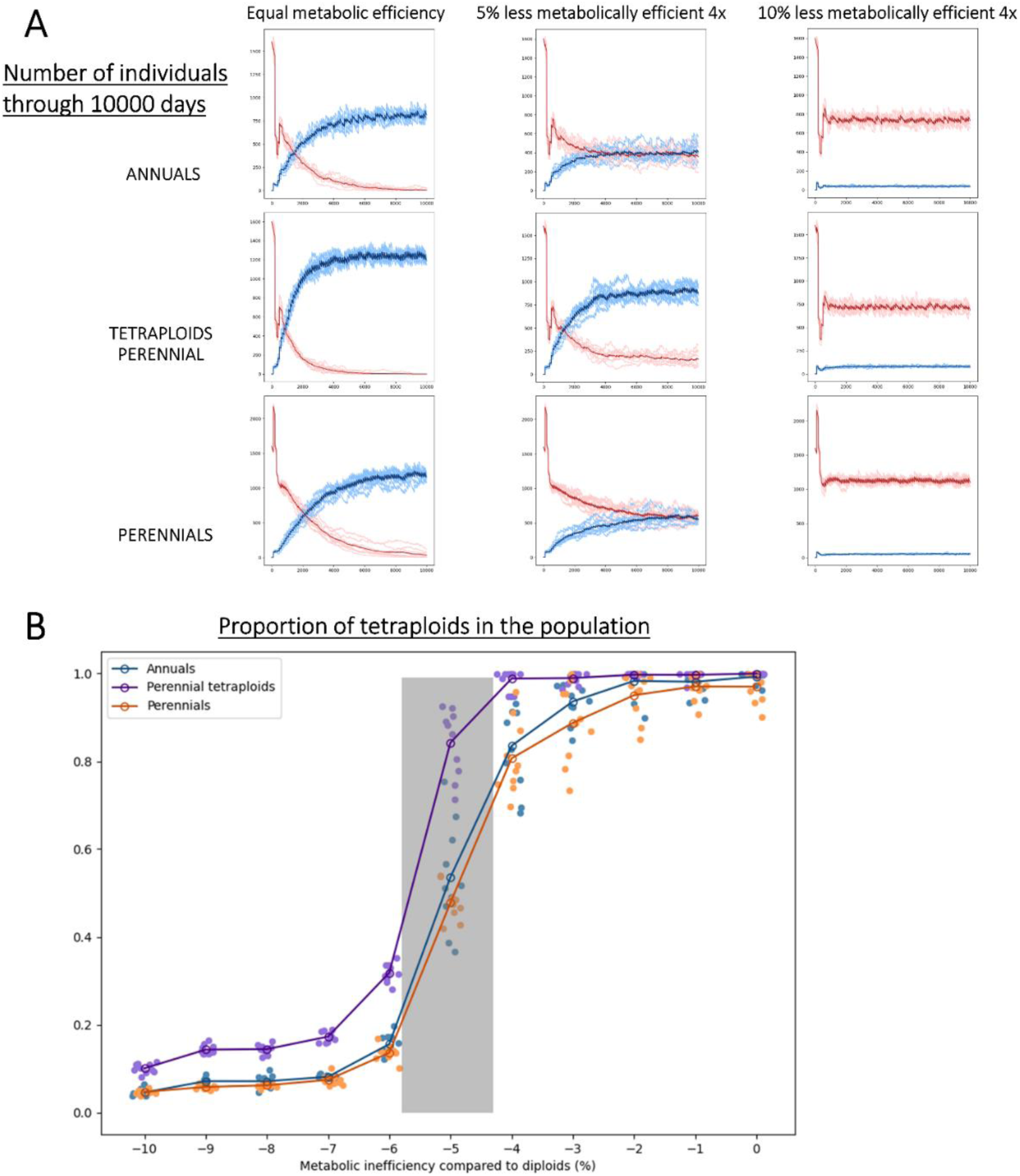
Main results of model simulations when polyploid formation is not drawn from Beta distribution with mean of 4.76% B(1, 20) but with a constant 4.76%. A) shows the number of individuals through 10,000 days, across simulated scenarios and metabolic efficiencies. Colors are red for diploids and blue for tetraploids. Rows correspond to the 3 life history scenarios: top row to the annuals scenario, bottom row to the perennials scenario, and middle to the scenario where only tetraploids exhibit perenniality. Columns represent tested metabolic efficiencies, with the first column showing results when tetraploid have an equal metabolic efficiency as diploids, and the second and third columns are showing results when tetraploids are 5 and 10% less efficient than diploids, respectively. Notice that these plots illustrate the mean of independent simulation in darker color shades of red and blue, while the lighter shades are all the independent simulations. B) Proportion of tetraploid individuals across metabolic inefficiencies of tetraploids in comparison to diploids. Results are shown for the three simulated scenarios: all individuals annual in blue, all individuals perennial in orange and tetraploid perennials in purple. Independent simulation runs are represented by scattered dots, and darker shades are the means. Notice the grey shaded area on the plot which represents the zone where coexistence is observed.

**Figure S5.**
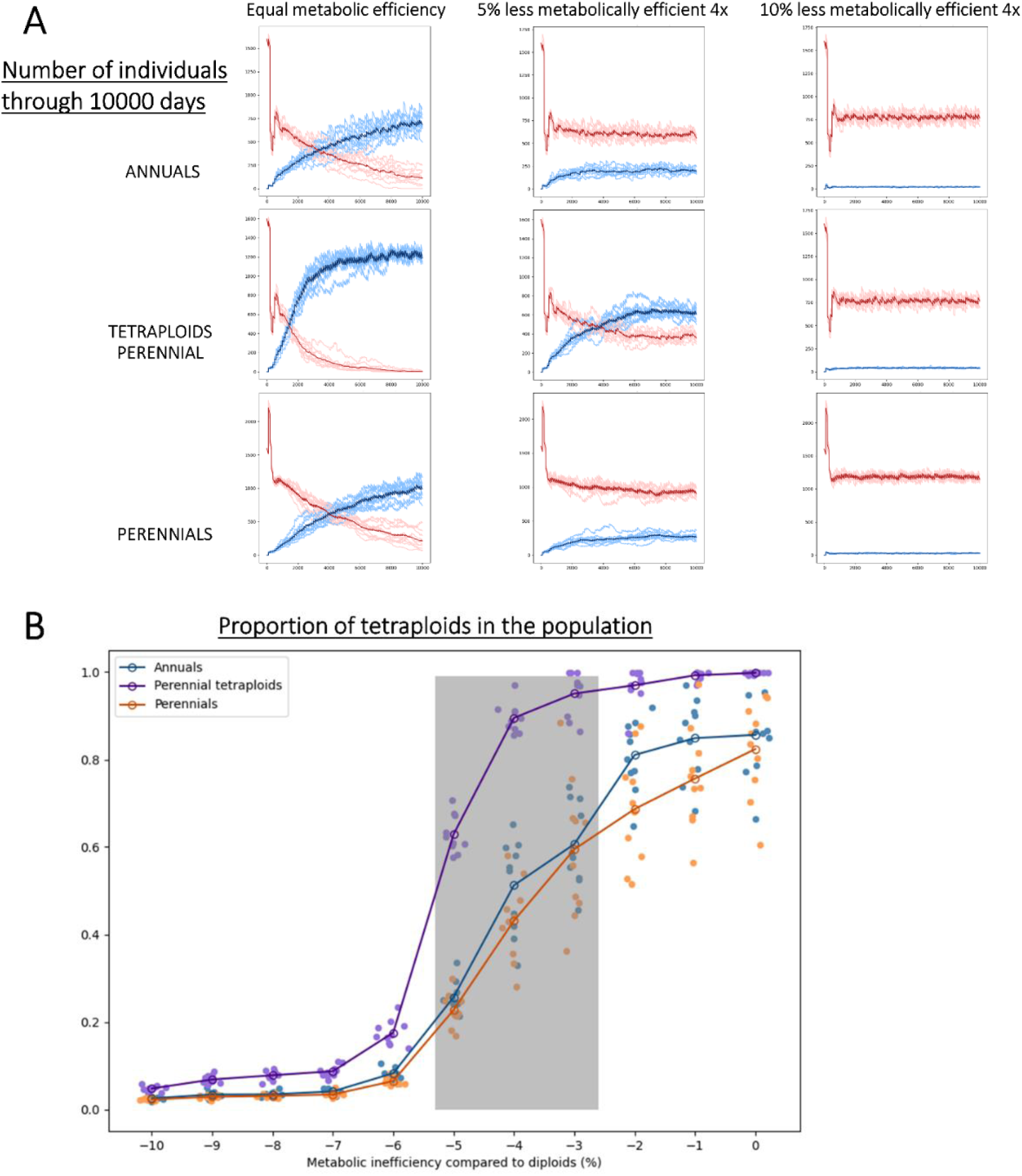
Main results of model simulations when polyploid formation is drawn from Beta distribution with mean of 2.38% B(2, 82). A) shows the number of individuals through 10,000 days, across simulated scenarios and metabolic efficiencies. Colors are red for diploids and blue for tetraploids. Rows correspond to the 3 life history scenarios: top row to the annuals scenario, bottom row to the perennials scenario, and middle to the scenario where only tetraploids exhibit perenniality. Columns represent tested metabolic efficiencies, with the first column showing results when tetraploid have an equal metabolic efficiency as diploids, and the second and third columns are showing results when tetraploids are 5 and 10% less efficient than diploids, respectively. Notice that these plots illustrate the mean of independent simulation in darker color shades of red and blue, while the lighter shades are all the independent simulations. B) Proportion of tetraploid individuals across metabolic inefficiencies of tetraploids in comparison to diploids. Results are shown for the three simulated scenarios: all individuals annual in blue, all individuals perennial in orange and tetraploid perennials in purple. Independent simulation runs are represented by scattered dots, and darker shades are the means. Notice the grey shaded area on the plot which represents the zone where coexistence is observed.

**Figure S6.**
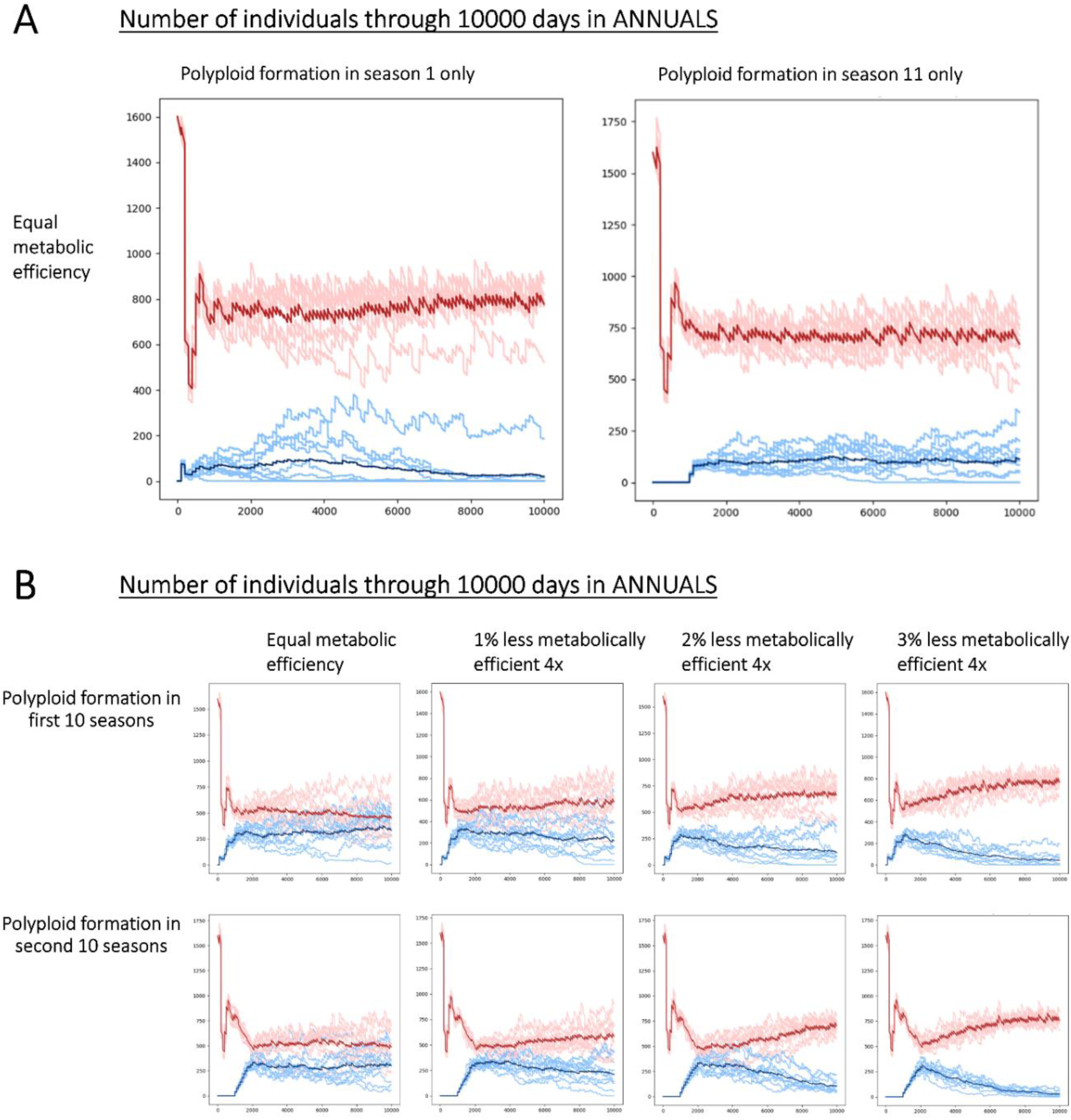
Number of individuals through 10,000 days, across scenario where all individuals are annuals and polyploid formation is not recurrent or constant. Color of lines are differentiating between diploids (depicted in red) and tetraploids (depicted in blue). Notice that these plots illustrate the mean of independent simulation in darker color shades of red and blue, while the lighter shades are all the independent simulations. A) shows the number of individuals when polyploids are equally metabolically efficient as diploids, but polyploid formation is allowed only during one season, either the first simulated season (left) or the eleventh season when the diploid population reaches carrying capacity (right). B) shows number of individuals when polyploid formation is allowed only during 10 season, either the first 10 seasons (top) or the second 10 seasons when the diploid population reached carrying capacity (bottom). The different columns represent tested metabolic efficiencies, with the first column showing results when tetraploid have an equal metabolic efficiency as diploids, and towards the right are lower efficiencies of polyploids one by one percent less than in diploids.

**Figure S7.**
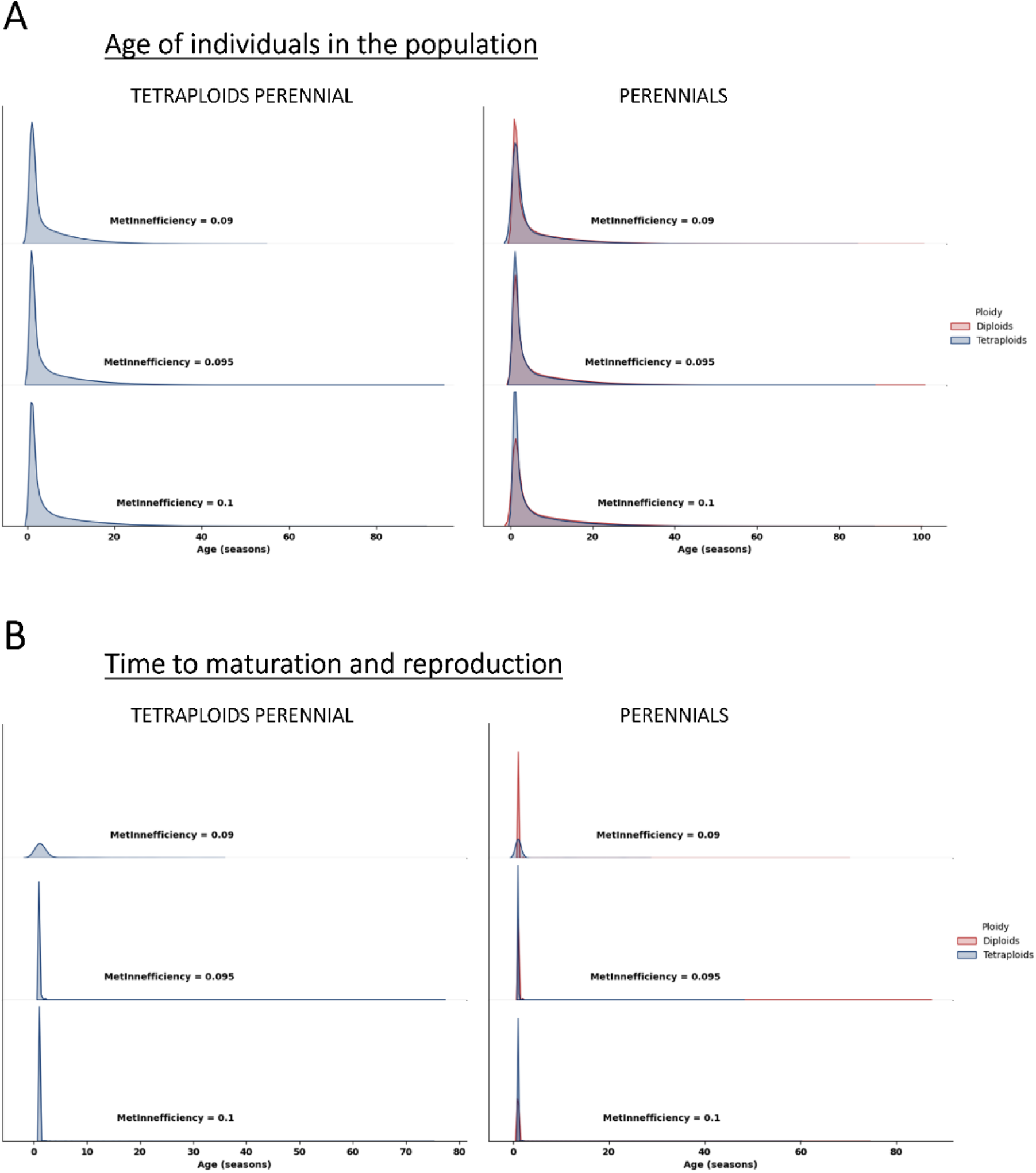
Distribution of ages of diploid and tetraploid individuals across simulated metabolic inefficiencies of tetraploids and life history strategies. A) shows the age distributions of all individuals, whereas B) shows age distributions of only the individuals which matured and successfully reproduced. Different columns correspond to the tested life history scenarios, with the left corresponding to the scenario where only tetraploids exhibit perenniality, and the right to the perennials scenario. Different rows correspond to the tested metabolic inefficiencies of tetraploids in comparison to diploids, with the top and middle rows where tetraploids are 10% and 5% less efficient, respectively, and the bottom row where they are equal. Notice that red colors represent diploids and blue represent tetraploids, and that the scenario of annual life histories is omitted as all ages are 1 season.

